# Girdin regulates dendrite morphogenesis and cilium position in two specialized sensory neuron types in *C. elegans*

**DOI:** 10.1101/2020.10.20.347823

**Authors:** Inna Nechipurenko, Sofia Lavrentyeva, Piali Sengupta

**Affiliations:** Department of Biology, Brandeis University, Waltham, MA, USA; Department of Biology and Biotechnology, Worcester Polytechnic Institute, Worcester, MA, USA

**Keywords:** Cilia, Girdin/GIV, Dendrite, Sensory neurons

## Abstract

Primary cilia are located at the dendritic tips of sensory neurons and house the molecular machinery necessary for detection and transduction of sensory stimuli. The mechanisms that coordinate dendrite extension with cilium position during sensory neuron development are not well understood. Here, we show that GRDN-1, the *Caenorhabditis elegans* ortholog of the highly conserved scaffold and signaling protein Girdin/GIV, regulates both cilium position and dendrite extension in the postembryonic AQR and PQR gas-sensing neurons. Mutations in *grdn-1* disrupt dendrite outgrowth and mislocalize cilia to the soma or proximal axonal segments in AQR, and to a lesser extent, in PQR. GRDN-1 is localized to the basal body and regulates localization of HMR-1/Cadherin to the distal AQR dendrite. However, loss of HMR-1 and/or SAX-7/LICAM, molecules previously implicated in sensory dendrite development in *C. elegans*, do not alter AQR dendrite morphology or cilium position. We demonstrate that GRDN-1 localization in AQR is regulated by UNC-116/Kinesin-1, and that correspondingly, *unc-116* mutants exhibit severe AQR dendrite outgrowth and cilium positioning defects. In contrast, GRDN-1 and cilium localization in PQR is modulated by LIN-44/Wnt signaling. Together, these findings identify upstream regulators of GRDN-1, and describe new cellspecific roles for this multifunctional protein in sensory dendrite development.

## INTRODUCTION

Neurons are highly polarized cells. Axons and dendrites house unique sets of structural and signal transduction proteins and exhibit distinct signaling and integration properties (Bentley and Banker, 2016; Takano et al., 2015). Sensory neurons represent a particularly intriguing example of neuronal polarization. The typically asynaptic dendrites in a subset of these neurons such as invertebrate chemo- and mechanosensory neurons and vertebrate photoreceptors and olfactory neurons contain specialized cilia at their distal ends (Keil, 1997; Menco, 1997; Rohlich, 1975; Ward et al., 1975). Cilia are evolutionarily conserved signaling organelles comprised of a microtubule-based axoneme core surrounded by a specialized membrane enriched in signaling molecules. Cilia are nucleated by a centriole-derived basal body that migrates toward the cell surface and docks at the plasma membrane; thus, the location of a cilium is necessarily dictated by the position of the basal body (Dawe et al., 2007; Elric and Etienne-Manneville, 2014; Tang and Marshall, 2012). How the location and anchoring of the basal body at the distal dendrite is coordinated with dendrite outgrowth during sensory neuron differentiation is not fully understood.

The *C. elegans* hermaphrodite contains sixty ciliated sensory neurons (Perkins et al., 1986; Ward et al., 1975). The majority of these neurons are born and differentiate in the embryo, with the exception of three types including the gas-sensing AQR and PQR neurons, which are generated postembryonically (Gray et al., 2004; Sulston and Horvitz, 1977; Sulston et al., 1983). Dendrites of sensory neurons extend anteriorly or posteriorly in the head and tail, respectively, and contain morphologically distinct cilia at their distal ends (Doroquez et al., 2014; Perkins et al., 1986; Ward et al., 1975). Sensory cilia are exposed either directly or indirectly to the external environment, or in the case of AQR and PQR, to the internal pseudocoelomic body fluid (Perkins et al., 1986; Ward et al., 1975; White et al., 1986). Since dendrite extension and ciliogenesis in the majority of ciliated sensory neurons occur during late embryonic development (Heiman and Shaham, 2009; Nechipurenko et al., 2017; Serwas et al., 2017), these processes are challenging to visualize *in vivo*. In contrast, AQR and PQR extend their dendrites during larval stages, allowing for investigations into the mechanisms underlying coordination of basal body localization and dendrite morphogenesis.

The multifunctional molecule GRDN-1/Girdin has recently been implicated in regulating basal body positioning and dendrite morphogenesis in *C. elegans* head amphid and URX/BAG sensory neurons, respectively (Cebul et al., 2020; Nechipurenko et al., 2016). The twelve pairs of sensory neurons in the bilateral amphid sense organs are born near the presumptive nose in the early embryo (Sulston et al., 1983). Following initial anterograde dendritic extension toward the developing nose (Fan et al., 2019), these dendrites ‘stretch’ from their attachment points at the nose as the cell bodies migrate posteriorly to their final positions (Heiman and Shaham, 2009; Low et al., 2019). Centrioles/basal bodies can be visualized in close proximity to the plasma membrane at the embryonic stages when amphid sensory dendrites begin to form (Nechipurenko et al., 2017; Serwas et al., 2017), indicating that in these neurons, basal bodies are likely anchored at the dendritic ends before the dendrites extend retrogradely. Similarly, the dendrites of the embryonically born URX and BAG sensory neurons extend retrogradely (Cebul et al., 2020), although localization of basal bodies has not been studied extensively in these neurons during development. In the amphid neurons, GRDN-1 acts cell autonomously to regulate basal body position and plays only a minor role in dendrite development (Nechipurenko et al., 2016). In contrast, in URX/BAG, GRDN-1 functions in the IL2so glia to regulate dendrite morphology; the role of this protein in ciliogenesis in these neurons remains to be determined (Cebul et al., 2020) (M. Heiman, personal communication). These observations suggest that dendrite extension and basal body positioning are regulated by partly independent mechanisms in different neuron types, and that Girdin contributes differentially to these processes in a cell type-specific manner.

In contrast to the retrograde dendrite extension exhibited by embryonic amphid, URX, and BAG sensory neurons, the dendrites of the postembryonic AQR and PQR neurons extend anterogradely in a manner similar to mammalian olfactory neurons (Kirszenblat et al., 2011; Li et al., 2017; McEwen et al., 2008; White et al., 1986). The distal dendritic end of PQR is ensheathed by the process of the PHso2L glial cell, while the AQR distal dendrite does not appear to be associated with glia (Hall and Russell, 1991; White et al., 1986). LIN-44/Wnt signaling from posterior hypodermal sources acts via the LIN-17/Frizzled receptor in PQR to define the site of dendrite emergence from the PQR cell body (Kirszenblat et al., 2011). Moreover, translocation of the basal body to the PQR dendritic tip has been shown to require dynein (Li et al., 2017). Additional mechanisms for AQR/PQR (referred to henceforth as A/PQR) dendrite morphogenesis, basal body localization, or ciliogenesis are unknown.

Here we show that Girdin acts via partly distinct pathways to regulate dendrite morphogenesis and basal body/cilium positioning in A/PQR. Girdin is localized to the basal body in both neuron types, and tracks the dendritic growth cone as the AQR dendrite extends anterogradely during postembryonic development. Loss of *grdn-1* results in mislocalization of basal bodies and cilia to ectopic subcellular locations in A/PQR. Moreover, A/PQR neurons exhibit a range of aberrant morphological phenotypes in *grdn-1* mutants. Localization of HMR-1 cadherin to the AQR distal dendrite is disrupted in *grdn-1* mutants; however, neuronal knockdown of HMR-1 alone, or together with loss of SAX-7/L1CAM function, does not affect AQR morphology or cilium position. We further demonstrate that UNC-116/Kinesin-1 and LIN-44/Wnt regulate GRDN-1 localization in AQR and PQR, respectively, and that both dendrite morphogenesis and cilium positioning in these neurons are compromised in the corresponding mutants. Taken together, our results describe a new cell type-specific role for Girdin in modulating dendrite morphology and cilium positioning in two specialized sensory neuron types, and further highlight the importance of cellular context in the regulation of of function of this versatile protein.

## MATERIALS AND METHODS

### *C. elegans* genetics

Animals were grown on nematode growth medium (NGM) at 20°C with *E. coli* OP50 bacteria as a food source. Transgenic animals carrying extrachromosomal arrays were generated by germline microinjection of plasmids at 10-50 ng/μl into wild-type animals. A transgenic line selected based on expression levels and transmission rate was subsequently crossed into the relevant mutant backgrounds for phenotypic analyses. For rescue experiments of *grdn-1*-dependent phenotypes, plasmids were injected in *grdn-1(hmn1)* mutant animals. At least two independent lines were examined for each extrachromosomal array generated in this work, and typically results from one representative line are reported. When necessary, *unc-122*p*::gfp*, *unc-122*p*::mcherry*, or *srg-47p::gfp* plasmids were used as co-injection markers (injected at 30-40 ng/μl). To knockout *unc-116* specifically in neurons, *tag-165p::Cre (nCre)* plasmid (Flavell et al., 2013) was injected together with a co-injection marker in the *unc-116(ce815)* background containing a floxed *unc-116* allele (Harterink et al., 2018). The presence of all mutations was confirmed by PCR and/or sequencing.

For experiments using larvae, eggs were collected either for 2 hours at room temperature, or overnight at 15°C, on standard NGM plates seeded with *E. coli* OP50 bacteria and allowed to develop for approximately 24 or 48 hours at 15°C prior to imaging. To obtain staged L1 animals, egg hatching was monitored continuously, and animals that hatched within a 30 min period were transferred to a separate plate and allowed to develop for 6-8 hours at 25°C prior to imaging. All strains used in this study are listed in Table S1.

### Molecular biology

The following promoters were used to drive expression in specific cells or tissues: *gcy-32p* (~0.7 kb, A/PQR and URX), *gcy-36p* (~1.0 kb, A/PQR and URX), *rab-3p* (~4.3 kb, gift from O. Hobert), and *grdn-1p* (~4.5 kb). cDNA sequences for *dyf-19, arl-13*, and *grdn-1* (isoform a) (Nechipurenko et al., 2016) and *xbx-1* (Kazatskaya et al., 2017) were subcloned under the relevant cell-specific promoters into a modified *C. elegans* expression plasmid pPD95.77 (a gift from A. Fire) or pMC10 (a gift from M. Colosimo) using standard cloning protocols. *snb-1* cDNA was amplified from a mixed-stage N2 cDNA pool, verified by sequencing, and subcloned under the relevant cell-specific promoter. To generate GRDN-1^ΔGCV^, a fragment of *grdn-1a* cDNA lacking sequences corresponding to the GCV amino acids was amplified by PCR and subcloned into the relevant expression vector to replace the corresponding full-length *grdn-1a* fragment. The sequence of the mutated *grdn-1* construct was verified by sequencing. All plasmids used in this work are listed in Table S2.

### CRISPR/Cas9-mediated homologous recombination

The auxin-inducible degron (AID) sequence was inserted immediately upstream of the *gfp* sequence in the LP172 *hmr-1(cp21[hmr-1::gfp + loxP])* I strain (Marston et al., 2016) using CRISPR/Cas9-mediated homologous recombination as described (Dokshin et al., 2018). To induce homologous recombination, an injection mixture comprised of the following reagents was injected into LP172 animals: *S. pyogenes* Cas9 Nuclease V3 (0.25 μg/μl, IDT), CRISPR-Cas9 tracrRNA (1 μM, IDT), *unc-122p::dsRed* (40 ng/μl); donor template (0.11 μg/μl, megamer ssDNA fragment, IDT) with 120-bp homology arms, crRNA (0.02μg/μl, IDT): rCrArArUrArUrUrUrCrUrGrUrCrGrUrCrArCrGrUrGrUrUrUrUrArGrArGrCrUrArUrGr CrU (r denotes RNA nucleotides). AID insertion was verified by PCR and sequencing.

### Auxin-inducible degradation of HMR-1

Transgenic strains containing *hmr-1::AID::gfp* and either *egl-17*p::*myr-mCherry* or *gcy-36*p::*myr-tagRfp* to label AQR (see Table S1) were injected with *gcy-32*p::*TIR1* plasmid and *unc-122*p::*dsRed* co-injection marker at 10 and 40 ng/μL, respectively. L4 hermaphrodites containing *hmr-1::AID::gfp, gcy-32*p::*TIR1* (selected on the basis of the *unc-122*p::*dsRed* co-injection marker), and *egl-17*p::*myr-mCherry* or *gcy-36*p::*myr-tagRfp* transgenes were placed on OP50-seeded NGM plates containing 2 mM auxin and allowed to age to young adults at 20°C overnight. Following a two-hour egg collection, adult animals were removed from auxin-containing plates, while eggs were allowed to develop at 15°C to L3 larvae or young adults, and AQR neurons from animals carrying all transgenes were imaged on a spinning disk confocal microscope.

### Microscopy

Larvae or one-day old adult hermaphrodites were immobilized in 10 mM tetramisole (Sigma) and mounted on 10% agarose pads set on microscope slides. Animals were imaged on an inverted spinning disk confocal microscope (Zeiss Axiovert with a Yokogawa CSU22 spinning disk confocal head). Complete *z*-stacks of A/PQR neurons were acquired at 0.27 μm intervals in Slidebook 6.0 (3i – Intelligent Imaging Innovations) using a Plan Apochromat 100X NA 1.4 or 63X NA 1.4 oil immersion objective. For experiments examining cilia position and GRDN-1 localization in the *lin-44* mutant background, adult or larval animals were imaged on an upright THUNDER Imager 3D Tissue (Leica). Complete *z*-stacks of PQR were acquired at 0.22 μm intervals in Leica Application Suite X software using an HC Plan Apochromat 63X NA 1.40-0.60 oil immersion objective. Acquired optical sections were rendered into maximum intensity projections using Slidebook 6.0 or Image J (NIH, Bethesda, MD). A/PQR morphology and localization of fluorophore-tagged proteins were quantified in Slidebook 6.0 or Image J.

### Statistical analysis

Prism 6 software (GraphPad, San Diego, CA) was used to carry out statistical analysis and generate all bar graphs and scatter plots.

## RESULTS

### GRDN-1 is localized to the basal body in AQR and PQR sensory neurons

Although the bipolar A/PQR neurons (Figure 1A) are known to be ciliated (Hall and Russell, 1991; White et al., 1986), cilia position and structure in these neurons have not been extensively characterized. In adult hermaphrodites, the cilia membrane-associated protein ARL-13::tagRFP was localized to the distal dendrites in both A/PQR (Figure 1B). Similar to observations in head sensory neurons, in which cilia emanate from the distal dendritic tips (Doroquez et al., 2014; Perkins et al., 1986; Ward et al., 1975), the cilium was present at the distal end of the PQR dendrite (Figure 1A-B) (Li et al., 2017). However, in AQR, the cilium was typically localized near, but not at, the distal dendritic tip (Figure 1A-B).

**Fig. 1.**
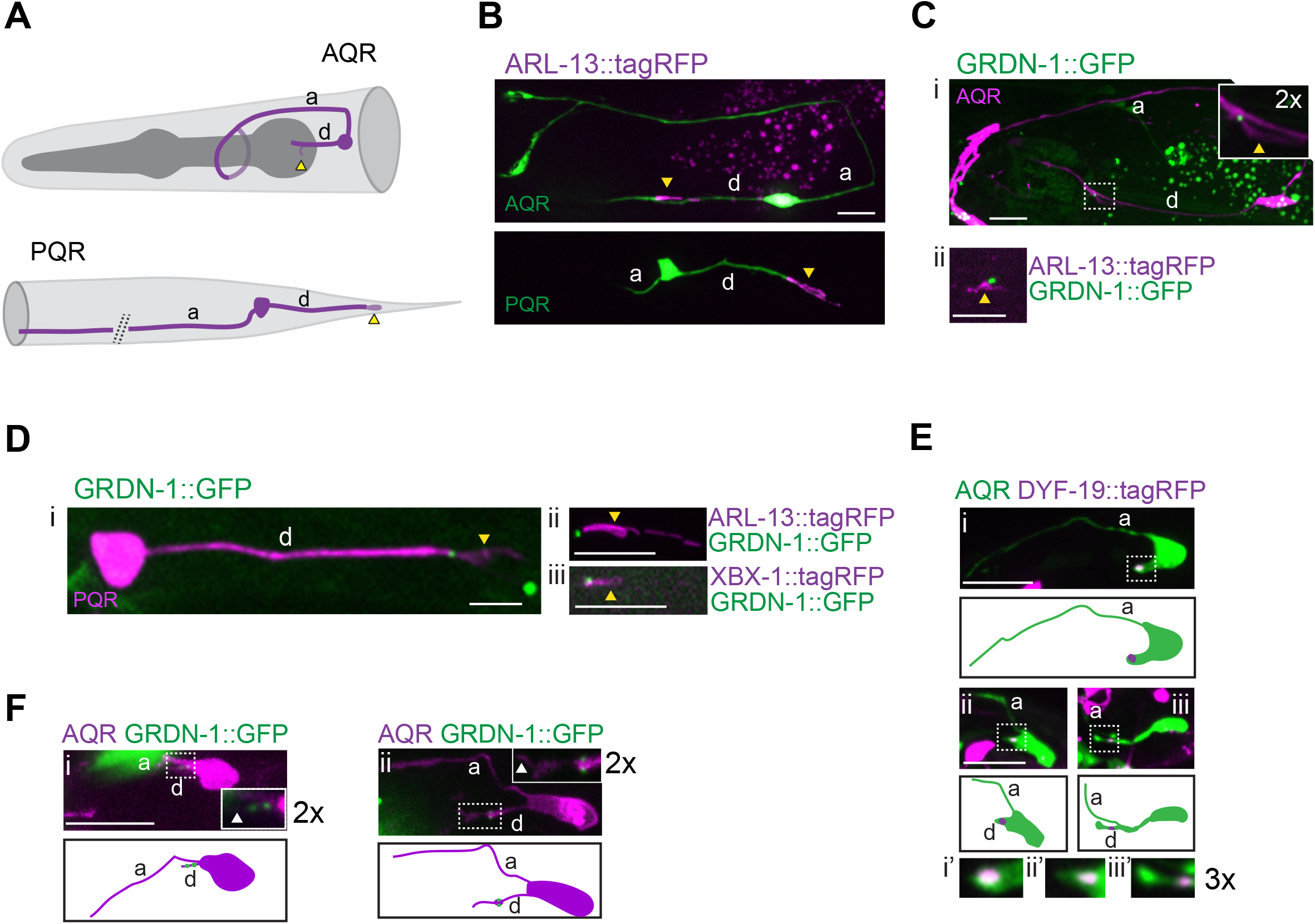
GRDN-1 is expressed in A/PQR and localizes to the cilia base in the distal dendrite. **(A)** Diagram of AQR (top) and PQR (bottom) neurons in an adult hermaphrodite. **(B)** Images of AQR (top) and PQR (bottom) in adults showing position of cilia labeled with *gcy-32p::ARL-13:*:tagRFP. A/PQR were visualized via expression of *gcy-37*p::GFP. **(C-D)** Images showing expression of GRDN-1::GFP in AQR (**Ci)** and PQR **(Di**), together with *gcy-32*p::ARL13::tagRFP in AQR (**Cii**) and PQR (**Dii**), and with *gcy-32*p::XBX-1::tagRFP in PQR (**Diii**). The cilium (dashed box) is enlarged (two-fold) in the inset in Ci. A/PQR were visualized with *gcy-32*p::mCherry. GRDN-1::GFP was expressed under *grdn-1* regulatory sequences in (**Ci**) and (**Di**) and *gcy-36* regulatory sequences in (**Cii)** and (**Dii-Diii**). All images are of adult hermaphrodites. **(E)** Images (**Ei-Eiii**, top) and cartoons (**Ei-Eiii**, bottom) showing localization of *gcy-36*p::DYF-19::tagRFP during the depicted stages of AQR dendrite outgrowth in L1 larvae. Dendritic tips (dashed boxes in **Ei-Eiii**, top) are enlarged three-fold in corresponding insets (**Ei’-Eiii’**). AQR was labeled with *gcy-37*p::GFP. **(F)** Images (**Fi-Fii**, top) and cartoons (**Fi-Fii**, bottom) of L1 larval AQR neurons marked with *egl-17*p::myr-mCherry expressing *gcy-36*p::GRDN-1::GFP. Insets show two-fold magnification of distal dendrites (white dashed boxes). Scale bars: 5 μm; anterior is at left. a - axon; d – dendrite; yellow arrowheads – cilium; white arrowheads −dendritic tips.

To investigate the expression and subcellular localization of GRDN-1 in A/PQR, we expressed the GFP-tagged full-length isoform a of GRDN-1 (Figure S1; henceforth referred to as GRDN-1::GFP) under its endogenous regulatory sequences. GRDN-1::GFP was expressed in A/PQR and localized to the cilia base in both neurons in adult hermaphrodites (Figure 1C-D). The dynein light intermediate chain protein XBX-1 localizes to the basal body and undergoes bidirectional movement along the ciliary axoneme (Schafer et al., 2003). The GRDN-1::GFP signal overlapped with the proximal-most region of the XBX-1::tagRFP signal in PQR (Figure 1Diii), suggesting that GRDN-1 localizes to the basal body in this neuron type.

AQR is born from the QR neuroblast in the L1 larval stage and completes migration to its final destination near the posterior bulb of the pharynx by ~9 hours posthatching (Middelkoop and Korswagen, 2014; Sulston and Horvitz, 1977). Upon reaching its final location, AQR initiates neuritogenesis (Chai et al., 2012). In contrast to observations in PQR (Kirszenblat et al., 2011; Li et al., 2017), outgrowth of the axon precedes that of the dendrite in AQR (Figure 1E). We noted that the basal body protein DYF-19::tagRFP was consistently associated with the distal segment of the extending dendrite, suggesting a coordination between dendrite outgrowth and basal body localization in AQR (Figure 1E). Similarly, GRDN-1::GFP puncta were found in proximity to the tip of the growing AQR dendrite (Figure 1F). These observations indicate that a basal body is associated with the distal dendrite during anterograde dendrite extension in AQR, and that Girdin is localized to the basal body in both A/PQR.

### GRDN-1 regulates basal body and cilia position in AQR

We asked whether GRDN-1 is required for correct basal body positioning in A/PQR as is the case in amphid sensory neurons (Nechipurenko et al., 2016). While the putative null mutation in *grdn-1* results in embryonic lethality (Nechipurenko et al., 2016), we had previously reported basal body positioning defects in amphid neurons in animals carrying viable *grdn-1(hmn1)* and *grdn-1(ns303)* hypomorphic alleles (Nechipurenko et al., 2016). The *ns303* mutation is an early stop codon that is predicted to produce a single isoform lacking the N-terminal putative microtubule-binding Hook domain, whereas the *hmn1* mutation is a late stop codon predicted to generate three truncated isoforms lacking a conserved PDZ binding motif and most of the predicted Gα-binding domain (Nechipurenko et al., 2016) (Figure S1). To quantify basal body localization in AQR, we scored DYF-19::tagRFP puncta as being present in the distal one-third of the dendrites, more proximal dendritic segments, or mislocalized to the cell bodies or axons. In wild-type AQR, DYF-19::tagRFP was found either within the distal one-third of the dendrite or in the dendritic regions slightly more proximal to the cell bodies in ~98% of neurons (Figure 2A-B). No localization was observed in soma or axons. In contrast, DYF-19::tagRFP was mislocalized to the AQR soma or proximal axon segments in a subset of both *grdn-1(ns303)* and *grdn-1(hmn1)* mutant animals, and was undetectable in an additional fraction of examined animals (Figure 2A-B).

**Fig. 2.**
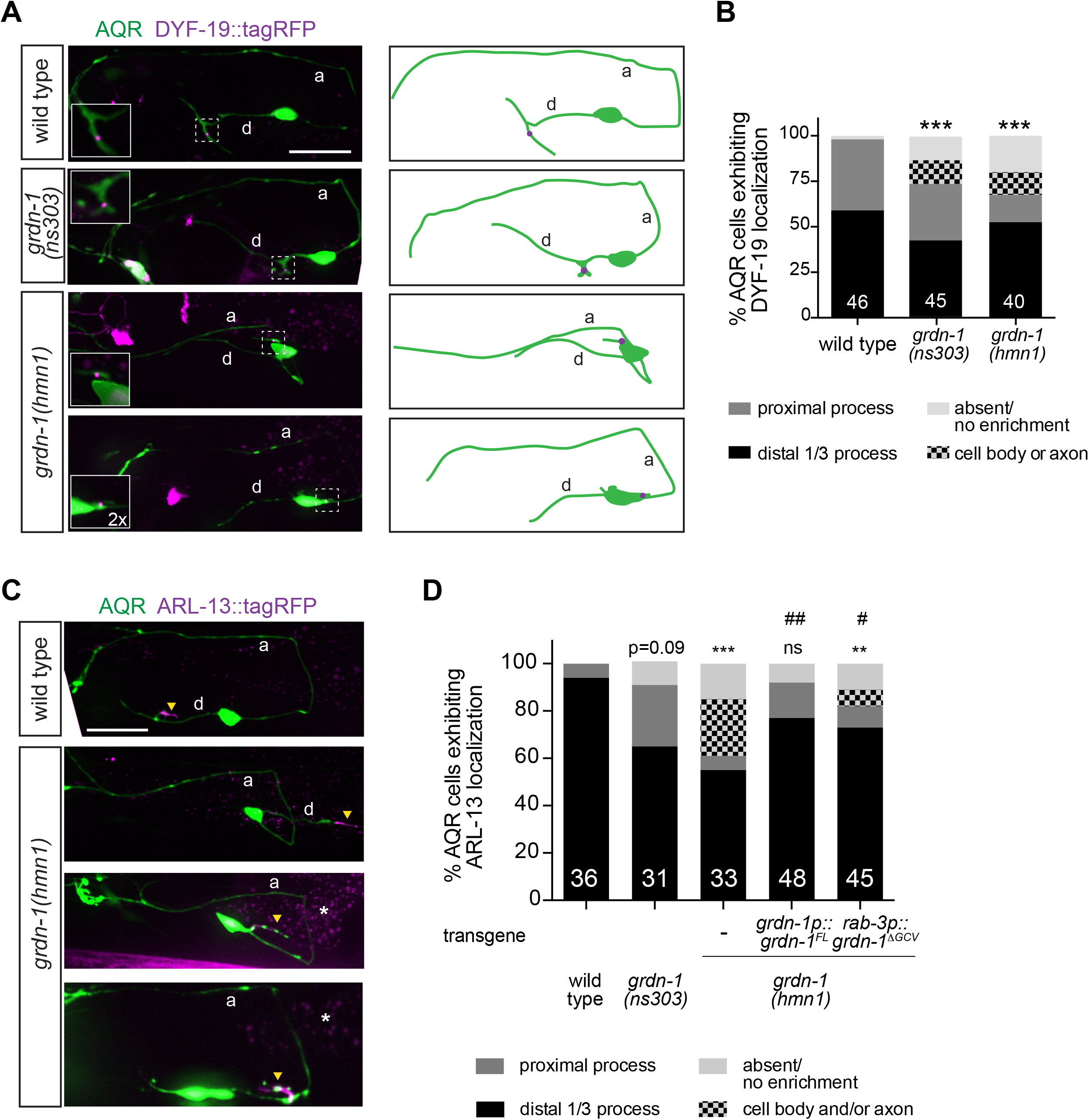
GRDN-1 regulates cilium position in AQR. **(A)** Images (left) and cartoons (right) showing localization of the basal body marker *gcy-36*p::DYF-19::tagRFP in AQR neurons of adult wild-type and *grdn-1* mutant animals. AQR was visualized with *gcy-37*p::GFP. Insets show two-fold magnification views of basal body regions (white dashed boxes). **(B)** Quantification of DYF-19::tagRFP localization in adult AQR neurons in the indicated genetic backgrounds. *** indicates different from wild type at p<0.001 (Fisher’s exact test). **(C-D)** Images **(C)** and quantification **(D)** of *gcy-32*p::ARL-13::tagRFP localization in AQR neurons in animals of the indicated genotypes. AQR was visualized with *gcy-37*p::GFP. Yellow arrowheads and asterisks mark cilia and gut autofluorescence, respectively. ** and *** indicate different from wild type at p<0.01 and p<0.001 (or the shown p-value); ns – not significant; # and ## indicate different from *grdn-1(hmn1)* at p<0.05 and p<0.01, respectively (Fisher’s exact test). In all image panels, a – axon; d – dendrite; anterior is at left. Scale bars: 10 μm. In all bar graphs, numbers indicate number of AQR neurons examined per genotype.

We noted that in *grdn-1* mutants, mislocalized DYF-19::tagRFP was occasionally detected at the base of a short cellular process, suggesting that cilia were being assembled at ectopic cellular locations in AQR (Figure 2A). Consistent with this hypothesis, the ciliary membrane protein ARL-13::tagRFP labeled cellular protrusions on the soma and proximal axonal segments of AQR in *grdn-1(hmn1)* mutants (Figure 2C-D). Although we did not detect similar ectopic structures in the somas/axons in *grdn-1(ns303)* animals, cilia were located in proximal dendritic segments of AQR in a larger percentage of *grdn-1(ns303)* mutants as compared to wild-type animals (Figure 2D). These findings suggest that basal body migration to the cell surface is largely unaffected in *grdn-1* mutants; instead, GRDN-1 may be required for correct basal body targeting to the distal sensory dendrite.

The ARL-13::tagRFP mislocalization phenotype in *grdn-1(hmn1)* animals was rescued upon expression of full-length wild-type *grdn-1a* cDNA under the endogenous *grdn-1* promoter (Figure 2D). The C-terminus of Girdin, which is deleted in the *grdn-1(hmn1)* allele, contains a three amino acid putative PDZ-binding motif (GCV) (Figure S1) (Enomoto et al., 2006; Oshita et al., 2003). We previously showed that deletion of C-terminal sequences including but not limited to the GCV motif resulted in mislocalization of GRDN-1 in amphid neurons (Nechipurenko et al., 2016). However, neuronal expression of GRDN-1 lacking the GCV motif alone (*grdn-1*^ΔGCV^) partially but significantly rescued the ARL-13::tagRFP mislocalization defects in *grdn-1(hmn1)* mutants (Figure 2D). We infer that GRDN-1 acts in neurons to regulate cilia positioning in AQR, and that the PDZ-binding GCV motif is largely dispensable for this process.

### GRDN-1 regulates AQR dendrite morphology

In the process of characterizing cilium position in adult hermaphrodites, we noted marked defects in AQR dendritic morphology in *grdn-1* mutants. Phenotypes included misrouted dendrites, which extended posteriorly; multipolar and unipolar, rather than bipolar, AQR morphology; dendrites and axons arising from a common process instead of opposites poles of the soma; as well as truncated dendrites (Figure 3A-B). While the penetrance of DYF-19::tagRFP localization defects were similar in both *grdn-1* alleles, dendritic defects were more penetrant in *grdn-1(hmn1)* than *grdn-1(ns303)* mutants (Figure 3B). Gross axon morphology was not altered in either mutant background. The dendritic defects in *grdn-1(hmn1)* mutants were partly but significantly rescued upon expression of a wild-type *grdn-1a* cDNA under the endogenous *grdn-1* or the pan-neuronal *rab-3* promoter (Figure 3B-C). Although the C-terminal GCV motif in GRDN-1 was shown to be important for retrograde extension of dendrites in the gas-sensing URX neurons (Cebul et al., 2020), a GRDN-1 construct lacking this motif partly but significantly rescued AQR dendrite morphology in *grdn-1(hmn1)* mutants (Figure 3B-C). These results indicate that GRDN-1 is required for dendritic morphogenesis in AQR, but unlike in URX, the GCV motif is not essential for GRDN-1 function in this neuron type.

**Fig. 3.**
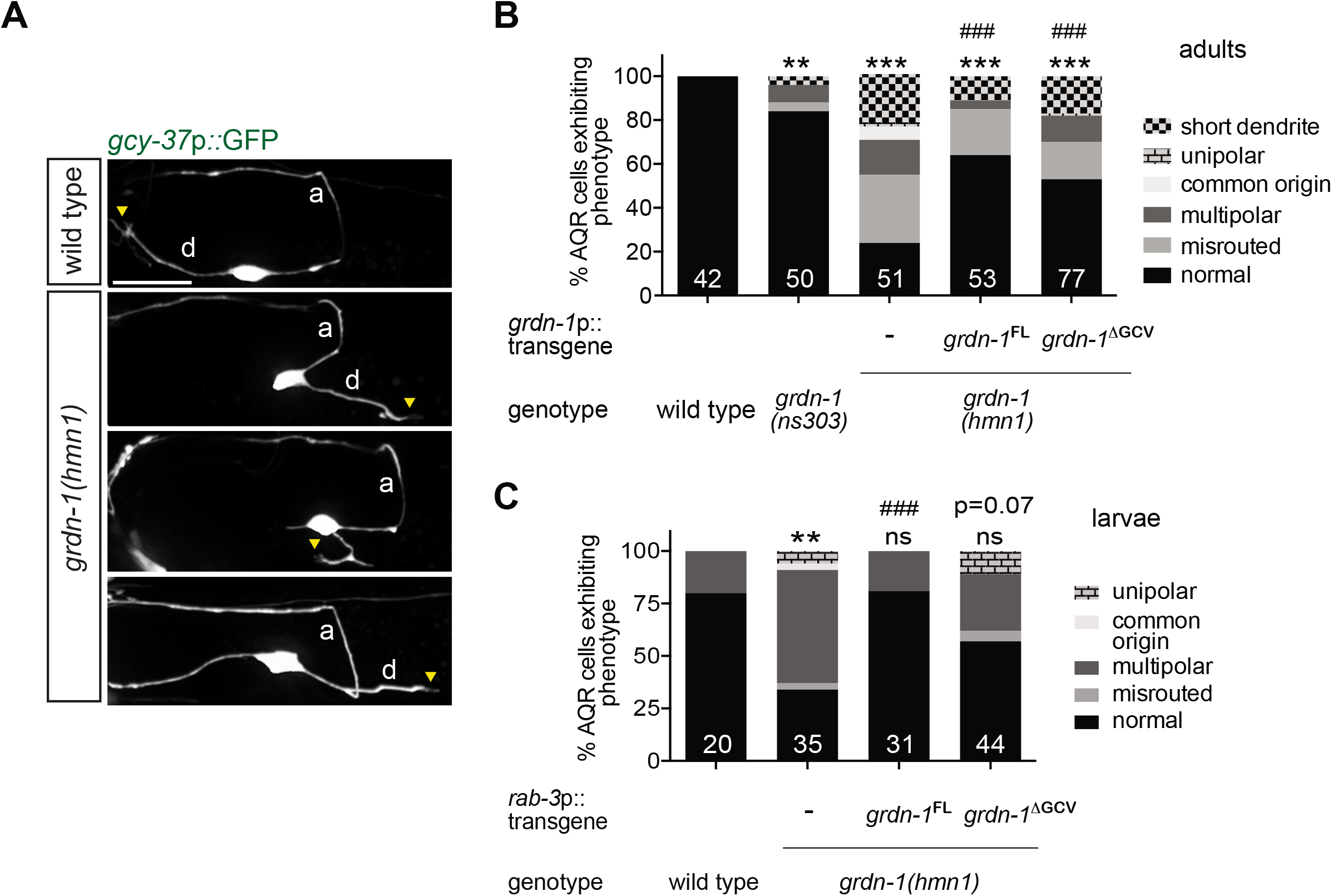
GRDN-1 is required for AQR dendrite development. Images **(A)** and quantification **(B, C)** of morphological defects in adult (**A, B**) and L2-L3 larval (**C**) AQR neurons of the indicated genotypes. Yellow arrowheads mark presumptive cilia; a – axon; d – dendrite; anterior is at left. Scale bar: 10 μm. FL – fulllength; ΔGCV – GCV motif deleted. ** and *** indicate different from wild type at p<0.01 and 0.001, respectively; ns - not significant; ### indicates different from *grdn-1(hmn1)* at p<0.001 or the indicated p-value; ns - not significant (Fisher’s exact test). Numbers indicate number of AQR neurons examined per genotype.

The presence of multiple neurites and reversed direction of dendrite outgrowth in *grdn-1* mutant AQR could arise from gross defects in neuronal polarity (Bentley and Banker, 2016; Yogev and Shen, 2017). To test this notion, we examined localization of the synaptic protein SNB-1::TagRFP in AQR of wild-type and *grdn-1(hmn1)* mutants. We noted that regardless of neuronal morphology, SNB-1::TagRFP was restricted to the synaptic regions in AQR axons of both wild-type and *grdn-1(hmn1)* animals (Figure S2). We conclude that it is unlikely that defects in AQR morphology observed in *grdn-1* mutants are due to global changes in neuronal polarity.

Since the basal body and cilium are associated with the distal end of the AQR dendrite, we asked whether cilium/basal body mislocalization and dendritic defects were correlated in *grdn-1* mutants. We scored cilium position as ‘wild-type’ when ARL-13::tagRFP was localized to the distal segment of a presumed dendrite, regardless of the direction of dendrite outgrowth. We found that in *grdn-1(hmn1)* mutants, ARL-13::tagRFP was present in the soma or proximal axonal segments in 50% of examined AQR neurons that exhibited altered morphology [n=28; *grdn-1(ns303)* animals were not examined due to the low penetrance of the AQR dendritic defect in this mutant background]. Conversely, we observed ectopic ARL-13::tagRFP localization to the soma or axon in 33% of morphologically wild-type AQR neurons in *grdn-1(ns303)* mutants (n=27). Together with the differential effects of *grdn-1(hmn1)* and *grdn-1(ns303)* mutations on basal body positioning and neuronal morphology, these results imply that GRDN-1 plays at least partly independent roles in basal body positioning and dendrite morphogenesis in AQR.

### Mutations in *grdn-1* affect cilia position and dendritic morphology in PQR

Although the dendrites of both AQR and PQR extend anterogradely in early larval stages, a key anatomical difference between AQR and PQR is that the distal end of the PQR, but not the AQR, dendrite is ensheathed by the PHso2L glial cell process (Hall and Russell, 1991; White et al., 1986). Recently, GRDN-1 was shown to act cell non-autonomously to regulate retrograde extension of glia-associated URX dendrites (Cebul et al., 2020). We asked whether GRDN-1 is required for anterograde extension of the glia-associated dendrite, and/or cilium position, in PQR.

While ARL-13::tagRFP was present at the distal tip of the PQR dendrite in 100% of wild-type animals, in *grdn-1* mutants, ARL-13::tagRFP was mislocalized to the PQR cell body or proximal dendritic segments in a subset of examined animals (Figure 4A-B). As in AQR, the ARL-13::tagRFP mislocalization phenotype was more penetrant in *grdn-1(hmn1)* than *grdn-1(ns303)* mutants and was fully rescued upon GRDN-1^ΔGCV^ expression under the pan-neuronal *rab-3* promoter (Figure 4A-B). PQR also exhibited a range of dendritic morphological defects in *grdn-1(hmn1)* mutants that were qualitatively similar to, but less penetrant than, those observed in AQR (Figure 4C-D). We noted that PQR dendrites were significantly shorter in *grdn-1(hmn1)* mutants; this defect was also fully rescued by GRDN-1^ΔGCV^ expression under the *rab-3* promoter (Figure 4E). We conclude that GRDN-1 acts in neurons to regulate PQR cilium position and dendrite morphology, and the C-terminal GCV motif is not required for the examined phenotypes in this neuron type. Moreover, GRDN-1 appears to play a more prominent role in AQR than in PQR morphogenesis.

**Fig. 4.**
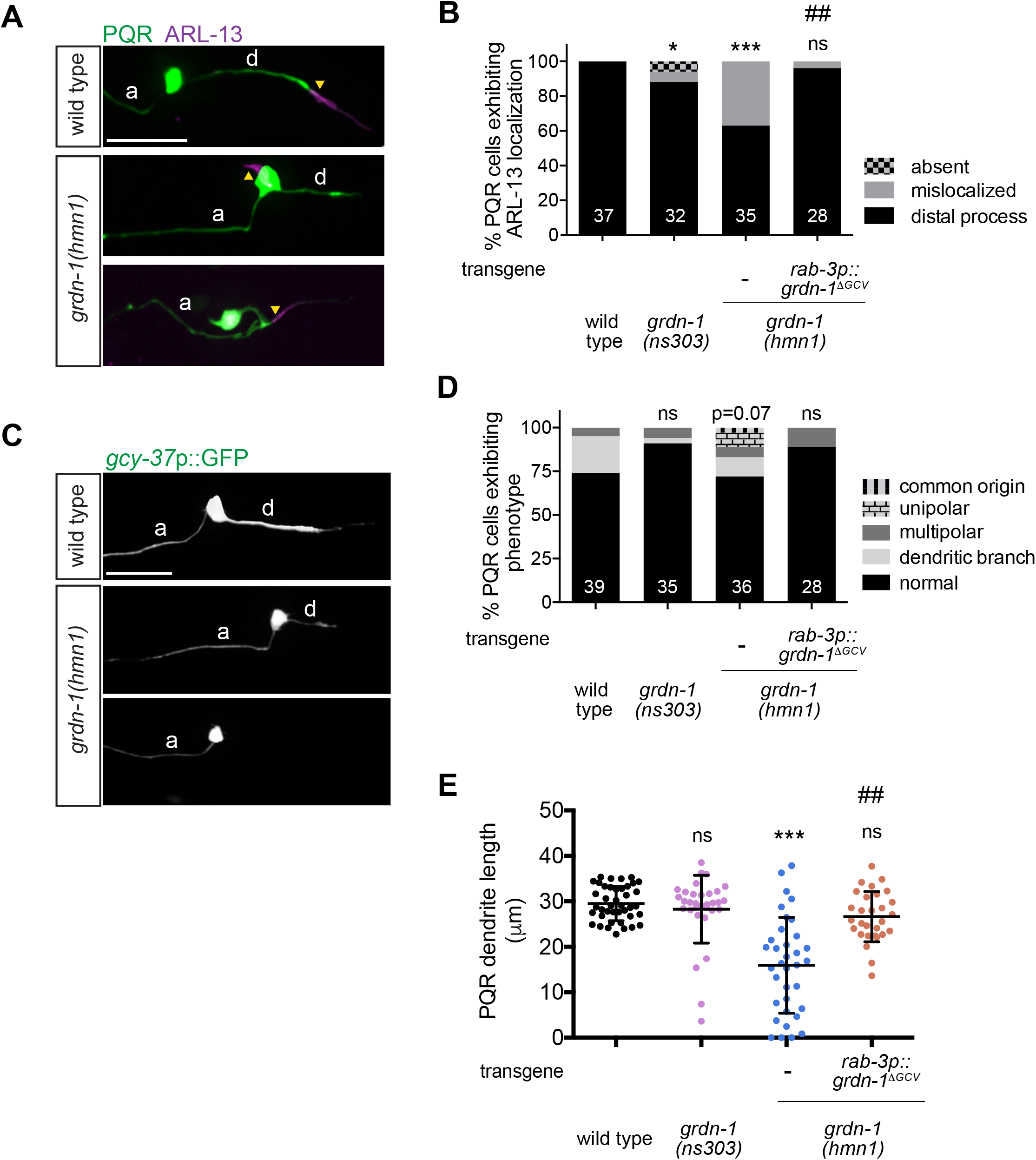
PQR morphology and cilium localization are differentially affected in *grdn-1* mutants. **(A–B)** Images **(A)** and quantification **(B)** of *gcy-32*p::ARL-13::tagRFP localization in PQR neurons of the indicated genotypes. PQR was visualized with *gcy-37*p::GFP. Yellow arrowheads mark cilia. * and *** indicate different from wild type at p<0.05 and p<0.001, respectively; ns – not significant; ## indicates different from *grdn-1(hmn1)* at p<0.01 (Fisher’s exact test). **(C–D)** Images **(C)** and quantification **(D)** of morphological defects exhibited by PQR neurons of the listed genotypes. p-value indicates different from wild-type; ns – not significant (Fisher’s exact test). **(E)** Quantification of PQR dendrite length in wild-type and *grdn-1* mutant animals. *** indicates different from wild type at p<0.001; ns – not significant (Kruskal-Wallis test with Dunn’s test for multiple comparisons). In all image panels, a – axon; d – dendrite; anterior is at left. Scale bars: 10 μm. In all bar graphs, numbers indicate number of PQR neurons examined per genotype.

### GRDN-1 regulates localization of HMR-1 cadherin, but not of SAX-7 L1CAM, in AQR

Multiple cell adhesion molecules mediate dendrite morphogenesis across species including in *C. elegans* (Berger-Muller and Suzuki, 2011; Dong et al., 2013; Lefebvre, 2017; McLachlan and Heiman, 2013; Salzberg et al., 2013; Seong et al., 2015). The HMR-1 classical cadherin, SAX-7 L1 cell adhesion molecule, DLG-1 Discs large homolog 1, and DYF-7 Zona Pellucida domain proteins are expressed in the embryonically born amphid sensory neurons in *C. elegans* and act redundantly to mediate the initial stages of dendrite extension in these cells (Fan et al., 2019). DYF-7 is also critical for retrograde extension of amphid, but not URX/BAG, sensory dendrites (Cebul et al., 2020; Heiman and Shaham, 2009; Low et al., 2019). Recently, GRDN-1 and SAX-7 were shown to mediate anchoring of URX/BAG sensory dendrites at the nose of the developing embryo (Cebul et al., 2020). Girdin also physically interacts with components of the cadherin-catenin complex during *Drosophila* embryogenesis (Houssin et al., 2015) and in human cancer cells (Wang et al., 2018), and regulates cadherin localization in some cellular contexts (Ichimiya et al., 2015; Muramatsu et al., 2015; Weng et al., 2014). We asked whether GRDN-1 acts via one or more of these identified cell adhesion molecules to regulate AQR morphology.

We first examined whether candidate cell adhesion molecules are expressed in the developing AQR neuron. We found that endogenously tagged HMR-1::GFP protein localized to the distal segment of the emerging AQR dendrite in wild-type larvae (Figure 5A), similar to its localization pattern at the distal tips of dendrites in embryonic amphid sensory neurons (Fan et al., 2019). Like DYF-19 and GRDN-1, HMR-1::GFP remained associated with the distal dendritic segment, as the AQR dendrite extended anteriorly (Figure 5A-B). We also observed localization of endogenously tagged SAX-7::GFP along the AQR dendrite, as well as in the soma and axon, of wild-type larvae (Figure S3A). However, DYF-7::sfGFP expressed under its endogenous promoter (Low et al., 2019) was detected in the AQR cell bodies and/or dendrite in only four out of 13 examined animals, suggesting that this expression may be an artifact of transgene overexpression (Figure S3B). We did not observe expression of endogenously tagged DLG-1::mNG protein in larval AQR (n=30).

**Fig. 5.**
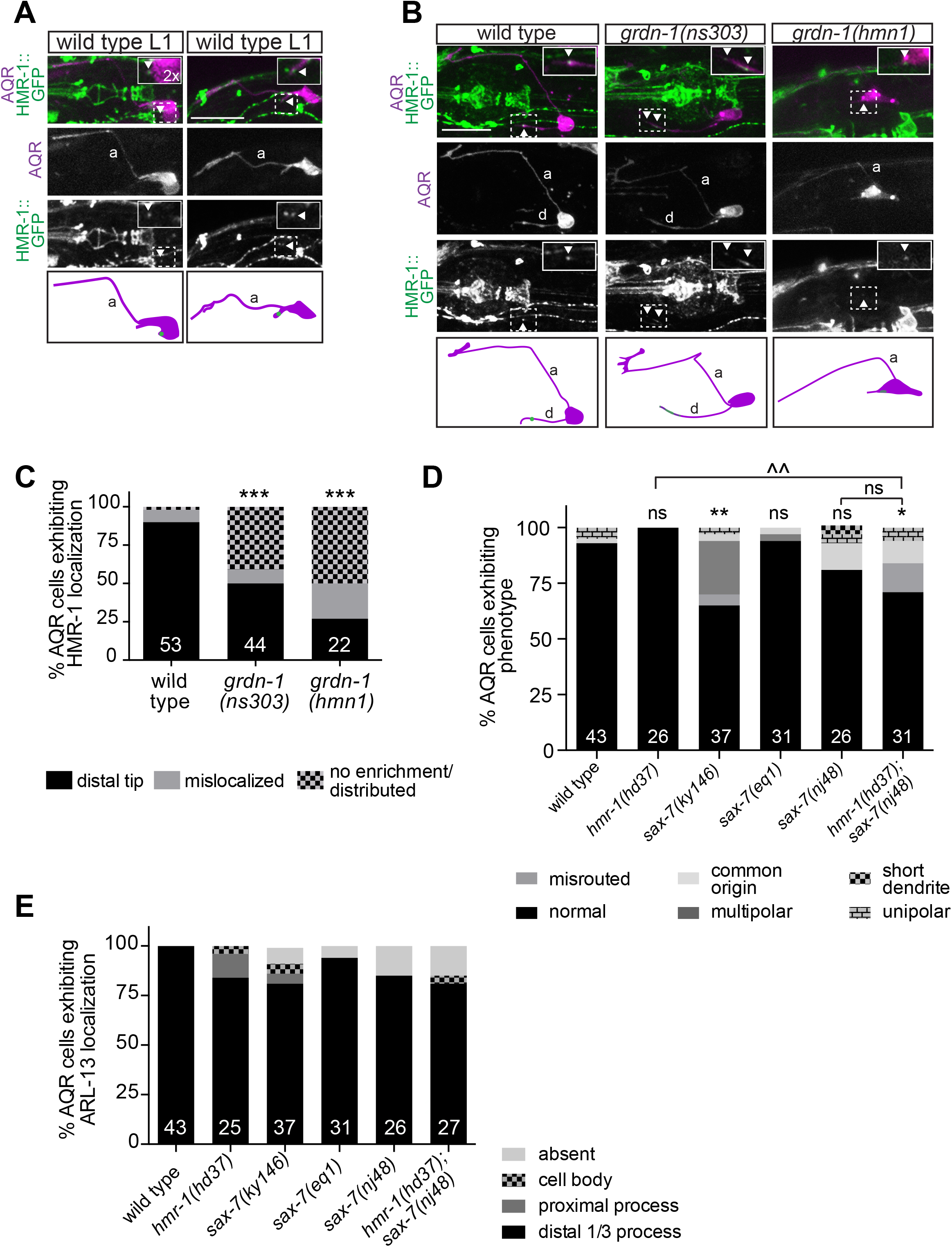
GRDN-1 regulates HMR-1 enrichment in AQR dendrites. **(A–B)** Images (top panels) and cartoons (bottom panels) showing localization of endogenously tagged HMR-1::GFP in AQR at different stages of dendrite outgrowth in wild-type L1 larvae **(A),** and in L2-L3 larvae of the indicated genotypes **(B)**. HMR-1::GFP localization in AQR (dashed boxes) is enlarged two-fold in all insets. AQR was visualized with *egl-17*p::myr-mCherry. White arrowheads mark HMR-1::GFP in AQR; a – axon; d – dendrite; anterior is at left. Scale bars: 5 μm. **(C–E)** Quantification of HMR-1::GFP localization in larvae **(C),** and AQR morphology (**D),** and *gcy-32*p::ARL-13::tagRFP localization **(E)** in adult animals of the indicated genotypes. In all bar graphs, numbers indicate number of AQR neurons examined per genotype. The phenotypes of the mutants indicated in **(E)** are not statistically different from that of wild type. *, ** and *** indicate different from wild type at p<0.05, p<0.01 and p<0.001, respectively; ^^ indicates different between indicated genotypes at p<0.01; ns – not significant (Fisher’s exact test).

We next asked whether GRDN-1 regulates AQR morphogenesis by modulating localization of HMR-1 and/or SAX-7. In both *grdn-1(hmn1)* and *grdn-1(ns303)* mutants, the HMR-1::GFP signal was no longer enriched in a punctum at the distal dendrite but instead was distributed, absent, or aberrantly localized to the AQR cell body (Figure 5B-C). HMR-1::GFP localization was similarly affected in both *grdn-1* mutant backgrounds (Figure 5C). In contrast, localization of SAX-7::GFP in AQR was unaffected in *grdn-1(hmn1)* mutants, and conversely, GRDN-1::GFP localized normally to the distal AQR dendrite in *sax-7(nj48)* animals (Figure S3A and S3C). We conclude that GRDN-1 is necessary for HMR-1, but not SAX-7, localization in AQR.

### Loss of HMR-1 and/or SAX-7 is not sufficient to alter AQR morphology or cilium positioning

Since GRDN-1 regulates HMR-1 localization, we investigated whether AQR morphogenesis or cilium positioning is altered in *hmr-1* mutants. Embryonic lethality of *hmr-1* null and *hmr-1*(RNAi) animals precludes examination of postembryonically generated AQR neurons in these genetic backgrounds (Costa et al., 1998). However, the *hmr-1(hd37)* allele is predicted to specifically disrupt the neuronal *hmr-1b* isoform and is viable (Steimel et al., 2010) (www.wormbase.org). Despite altered HMR-1::GFP localization in AQR in *grdn-1* mutants, we observed no significant defects in AQR morphology or cilia position in *hmr-1(hd37)* animals (Figure 5D-E). Reasoning that this hypomorphic mutation may not result in sufficient reduction of *hmr-1* function to cause defects in AQR, we depleted HMR-1 in a subset of sensory neurons including AQR using the auxin-inducible protein degradation (AID) system (Holland et al., 2012; Nishimura et al., 2009; Zhang et al., 2015). We introduced the degron sequence into the endogenous *gfp*-tagged *hmr-1* locus and generated strains expressing this fusion protein together with the TIR1 F-box protein driven under the *gcy-32* promoter that is transcriptionally active in A/PQR and URX (Yu et al., 1997). The HMR-1::GFP signal was no longer enriched at the distal tip of the AQR dendrites in the presence of auxin in L3 larvae grown on auxin-containing plates (Figure S4A). However, AQR morphology was largely unaffected in animals raised under these conditions (Figure S4B). Thus, although GRDN-1 is required for proper HMR-1 localization in AQR, neuron-specific reduction of HMR-1 function alone is not sufficient to alter AQR morphology or cilium position.

Since *sax-7* is also expressed in AQR and is known to function redundantly with *hmr-1* in several developmental contexts (Fan et al., 2019; Grana et al., 2010), we examined whether SAX-7 contributes to AQR morphogenesis. We observed weak but significant defects in AQR dendritic morphology in *sax-7(ky146)* but not in *sax-7(nj48)* or *sax-7(eq1)* animals (Figure 5D). Since the *sax-7(ky146)* allele has been reported to generate aberrant readthrough transcripts particularly in the nervous system (Wang et al., 2005), it is possible that the low penetrance AQR defects observed in this mutant background arise due to neomorphic SAX-7 function. AQR morphological defects in *sax-7(nj48)* mutants were not enhanced by additional loss of *hmr-1* (Figure 5D). Similarly, we did not observe significant defects in AQR cilium positioning in any of the examined *sax-7* alleles or in *hmr-1(hd37); sax-7(nj48)* double mutants (Figure 5E). Together, these results suggest that HMR-1 and SAX-7 are not required for dendrite outgrowth and cilium positioning in AQR, or that these proteins act redundantly with additional molecules to regulate these aspects of AQR development.

### UNC-116/Kinesin-1 regulates GRDN-1 localization in AQR

The mechanisms that regulate GRDN-1 function and localization in the context of sensory dendrite development or ciliogenesis are unknown. The KIF5a kinesin-1 motor was previously shown to regulate Girdin localization to the Golgi and cell adhesion contacts in mammalian cells (Muramatsu et al., 2015). We examined whether UNC-116/Kinesin-1 similarly regulates GRDN-1 localization in the AQR dendrite. We found that GRDN-1::GFP was localized to the soma in ~65% of examined *unc-116(e2310)* mutants, as compared to 12% of wild-type animals (Figure 6A-B). Consistent with a role for GRDN-1 in localizing HMR-1 to distal AQR dendrites, HMR-1::GFP puncta were also frequently mislocalized to the cell body in *unc-116(e2310)* animals (Figure 6C-D).

**Fig. 6.**
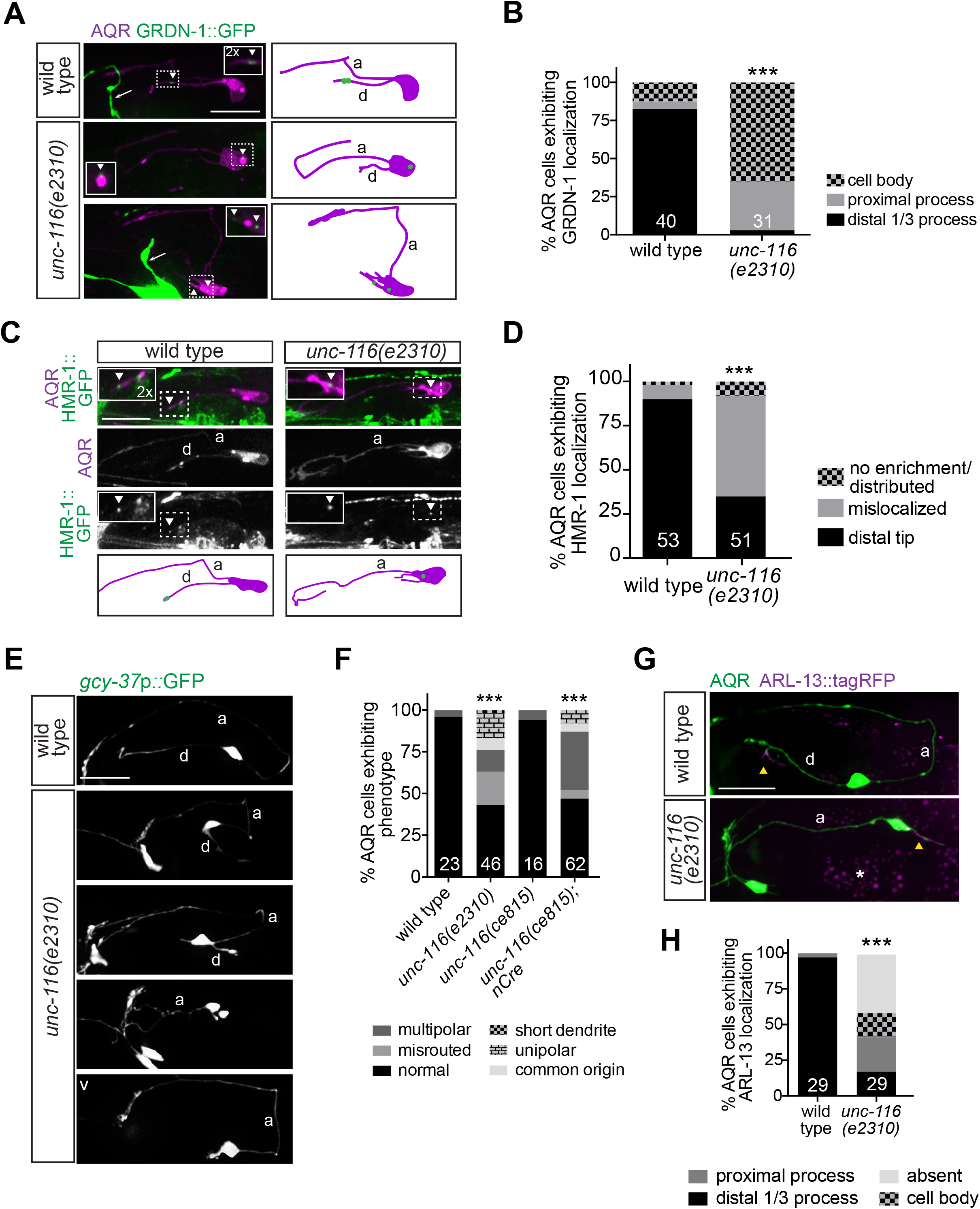
Kinesin-1/UNC-116 regulates GRDN-1 localization and AQR morphology. **(A)** Images (left) and cartoons (right) of *gcy-36*p::GRDN-1::GFP localization in AQR of wild-type and *unc-116* mutant larvae. AQR was labeled with *egl-17*p::myr-mCherry. GRDN-1::GFP signal in AQR (white dashed boxes) is magnified two-fold in all insets. White arrowheads mark GRDN-1::GFP; white arrows point to ASI sensory neuron labeled with *srg-47*p::GFP in the background. **(B)** Quantification of GRDN-1::GFP localization in wild-type and *unc-116* L2-L3 larvae. *** indicates different from wild type at p<0.001 (Fisher’s exact test). **(C)** Images (top panels) and cartoons (bottom panels) showing localization of the endogenously tagged HMR-1::GFP in wild-type and *unc-116* mutant L2-L3 larvae. HMR-1::GFP signal in AQR (white dashed boxes) is magnified two-fold in all insets and marked with white arrowheads. AQR was labeled with *egl-17*p::myr-mCherry. **(D**) Quantification of HMR-1::GFP localization in wild-type and *unc-116* L2-L3 larvae. Data for wild type are repeated from Figure 5C. *** indicates different from wild type at p<0.001 (Fisher’s exact test). (**E–F)** Images **(E)** and quantification **(F)** of AQR morphological phenotypes in the indicated genetic backgrounds. *** indicates different from wild type at p<0.001 (Fisher’s exact test). **(G–H)** Images **(G)** and quantification **(H)** of *gcy-32*p::ARL-13::tagRFP localization in AQR of adult wild-type and *unc-116* mutant animals. Yellow arrowheads and white asterisk mark cilia and gut autofluorescence, respectively. *** indicates different from wild type at p<0.001 (Fisher’s exact test). In all image panels, a – axon; d – dendrite; anterior is at left. Scale bars: 10 μm. In all bar graphs, numbers indicate number of AQR neurons examined per genotype.

Since UNC-116 regulates GRDN-1 localization in AQR, we asked whether *unc-116* mutants display defects in AQR morphology or cilium positioning similar to those observed in *grdn-1* mutant animals. Since null mutations in *unc-116* result in embryonic lethality (Byrd et al., 2001), we examined AQR phenotypes in animals carrying the hypomorphic *unc-116(e2310)* allele (Patel et al., 1993; Yan et al., 2013). As shown in Fig. 6E, *unc-116(e2310)* mutants exhibited a range of AQR dendritic phenotypes that resembled those observed in *grdn-1* mutants and included misrouted and truncated dendrites, as well as multipolar and unipolar neuronal morphologies. Moreover, animals in which *unc-116* was knocked out in neurons via Cre-loxP-mediated recombination (Harterink et al., 2018), exhibited AQR morphological defects similar to those in *unc-116(e2310)* mutants (Figure 6F). In addition to altered morphology, *unc-116(e2310)* mutant AQR neurons displayed ectopic localization of ARL-13::tagRFP (Figure 6G-H), with a subset of examined cells lacking expression possibly due to cilia loss (Figure 6H). In contrast, PQR neurons in *unc-116* mutants exhibited relatively weak dendritic and ciliary defects (Figure S5A-B and D-E). As in *grdn-1* mutants, PQR dendrites in *unc-116* animals were also significantly shorter compared to wild type (Figure S5C). We conclude that UNC-116 regulates GRDN-1 localization to the distal AQR dendrite, and is required for dendrite outgrowth and cilium positioning in AQR, and to a lesser extent, in PQR neurons.

### LIN-44/Wnt regulates GRDN-1 localization in PQR

We noted that the PQR dendrite defects in *grdn-1* mutants, although less penetrant, were qualitatively similar to those previously reported in *lin-44*/Wnt mutants (Kirszenblat et al., 2011). We hypothesized that GRDN-1 may mediate a subset of LIN-44-mediated functions in PQR morphogenesis. We first tested whether LIN-44 regulates GRDN-1 localization in PQR. While GRDN-1::GFP localized to the PQR distal dendrite in 97% of wild-type animals, only 20% of *lin-44(n1792)* mutants exhibited normal GRDN-1::GFP localization (Figure 7A-B). In the remaining fraction of *lin-44(n1792)* mutants, GRDN-1::GFP localized to ectopic cellular compartments including the soma and dendritic base, or was undetectable (Figure 7A-B), indicating that Wnt signaling plays a major role in regulating GRDN-1::GFP localization in PQR.

**Fig. 7.**
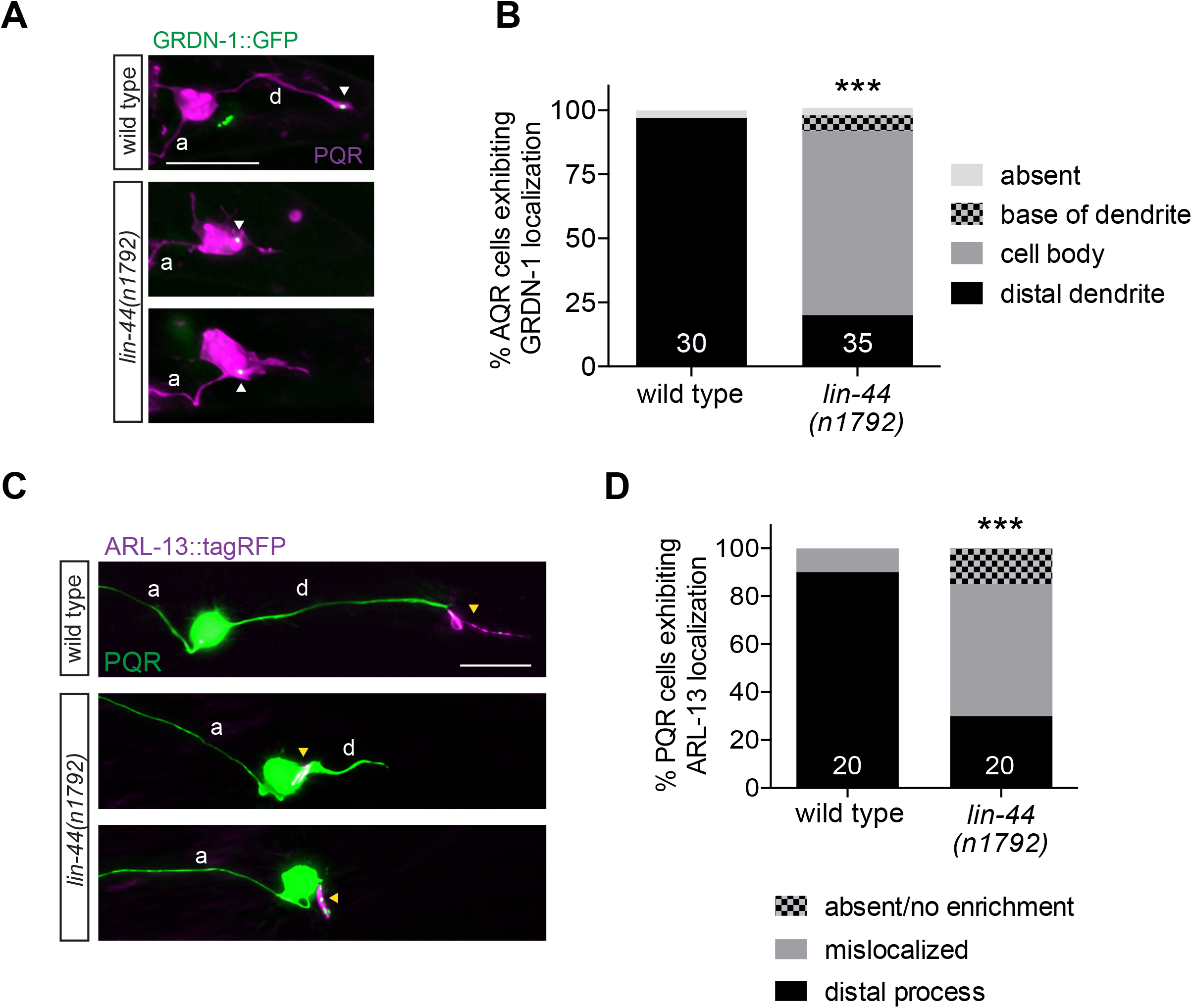
LIN-44/Wnt regulates GRDN-1 localization and cilium position in PQR. **(A-B)** Images **(A)** and quantification **(B)** of *gcy-36*p::GRDN-1::GFP localization in PQR of wild-type and *lin-44* mutant adult hermaphrodites. PQR was labeled with *gcy-37p::GFP*. White arrowheads indicate GRDN-1::GFP. *** indicates different from wild type at p<0.001 (Fisher’s exact test). **(C–D)** Images **(C)** and quantification **(D)** of *gcy-32*p::ARL-13::tagRFP localization in PQR of adult wild-type and *lin-44* mutant animals. Data for wild type are repeated from Figure S5E.Yellow arrowheads mark cilia. *** indicates different from wild type at p<0.001 (Fisher’s exact test). In all image panels, a – axon; d – dendrite; anterior is at left. Scale bars: 10 μm. In all bar graphs, numbers indicate number of PQR neurons examined per genotype.

A role for Wnt signaling in regulating cilium position in PQR has not been previously described. To test whether mislocalization of GRDN-1 in *lin-44* mutants is correlated with altered cilium positioning in PQR, we examined localization of ARL-13::tagRFP in this neuron type. We found that ARL-13::tagRFP localized normally to the PQR distal dendrite in only 30% of *lin-44(n1792)* mutants, and was present on the soma or diffusely distributed throughout the neuron in 70% of examined animals (Figure 7C-D). These results suggest that Wnt signaling regulates both dendritic morphology and cilium positioning in PQR, possibly in part via regulation of GRDN-1 localization.

## DISCUSSION

Here we report that GRDN-1 acts in neurons to regulate dendrite outgrowth and cilia position in the postembryonic AQR and PQR sensory neurons. In both neuron types, GRDN-1 is localized to the basal body. In AQR, the basal body is associated with the distal dendrite as it undergoes anterograde extension in early larvae. We find that in *grdn-1* mutants, the cilium in AQR is frequently mispositioned to the soma or proximal axonal segments, and dendrites exhibit a range of outgrowth defects. Similar defects in cilium localization and qualitatively similar but less penetrant defects in dendrite morphogenesis were also observed in PQR suggesting that GRDN-1 plays a more critical role in AQR development. We also show that GRDN-1 regulates HMR-1 but not SAX-7 localization in AQR, although loss of either molecule singly or together does not affect the examined aspects of AQR development. GRDN-1 localization is regulated by UNC-116/Kinesin-1 in AQR, and *unc-116* mutants in turn exhibit dendritic and cilium positioning phenotypes resembling those of *grdn-1* mutants. In contrast, GRDN-1 localization in PQR is regulated by LIN-44/Wnt signaling. Together, these results describe a new cell-specific role for Girdin and identify its upstream regulators in two ciliated sensory neuron types in *C. elegans*.

GRDN-1 may act via distinct mechanisms to regulate dendrite morphology and basal body/cilium positioning in AQR. This conclusion is based on the observations: 1) that the *grdn-1(ns303)* and *grdn-1(hmn1)* hypomorphic mutations result in qualitatively and quantitatively similar defects in basal body positioning in AQR, but only *grdn-1*(*hmn1*) animals exhibit severe defects in AQR dendrite outgrowth, and 2) that the dendrite outgrowth and basal body positioning defects appear to be uncorrelated in AQR in *grdn-1* mutants. GRDN-1 also acts in distinct pathways to regulate dendrite morphology and basal body positioning in other *C. elegans* sensory neuron types. In amphid sensory neurons, GRDN-1 regulates basal body position in part via CHE-10/Rootletin and the apical junction component AJM-1, but has a minor effect on dendrite anchoring (Nechipurenko et al., 2016). In URX/BAG, GRDN-1 acts in glia to regulate dendrite morphology, but it is unknown whether GRDN-1 also plays a role in regulating basal body/cilium position in these sensory neurons (Cebul et al., 2020). Although HMR-1::GFP was mislocalized in AQR in *grdn-1* mutants, we did not detect any defects in either AQR dendritic morphology or cilia positioning in *hmr-1* single or *hmr-1; sax-7* double mutants, indicating that the effectors of GRDN-1 in regulating different aspects of AQR development remain to be identified. Mammalian Girdin has been shown to act as a hub for multiple signaling pathways in different cellular contexts (Aznar et al., 2016; Garcia-Marcos et al., 2015). Since GRDN-1 is expressed in many neuronal and non-neuronal cell types in *C. elegans* (Nechipurenko et al., 2016), it will be interesting to determine the extent to which GRDN-1 function is modulated by diverse signaling pathways in different cell types.

We find that UNC-116/Kinesin-1 is required directly or indirectly for GRDN-1 localization to distal dendrites in AQR, whereas LIN-44/Wnt signaling regulates GRDN-1 localization in PQR. Reduction or loss of UNC-116 function results in aberrant neuronal morphology and cilium positioning in AQR. Similarly, mutations in *lin-44* lead to strong defects in PQR dendritic morphology (Kirszenblat et al., 2011), and we find that these mutants also exhibit strong defects in PQR cilium positioning. While AQR phenotypes are quantitatively similar in *unc-116* and *grdn-1(hmn1)* mutants, PQR phenotypes are more penetrant in *lin-44* than in *grdn-1* mutant animals. It is possible that this difference arises due to the hypomorphic nature of examined *grdn-1* alleles. Alternatively, GRDN-1 may function in parallel with other molecule(s) downstream of Wnt signaling to regulate PQR morphogenesis. A feature shared by amphid and PQR, but not AQR, neurons is that their dendrites are associated with glia (Doroquez et al., 2014; Hall and Russell, 1991; Perkins et al., 1986; Ward et al., 1975; White et al., 1986). Glial cells of the amphid organ secrete the DEX-1 tectorin-like molecule which acts with DYF-7 to facilitate anchoring of amphid dendritic tips at the nose (Heiman and Shaham, 2009). We speculate that GRDN-1-independent, but possibly LIN-44-dependent, glial interactions may similarly maintain PQR dendrite morphology upon reduction of *grdn-1* function. Indeed, mutations in *lin-44* affect development of the PHso2L glia (Herman and Horvitz, 1994; Herman et al., 1995), and ablation of these glia result in dendritic extension defects in PQR (Kirszenblat et al., 2011). Structural interactions with surrounding epidermal and muscle cells via multiple protein complexes have also been shown to play a major role in regulating the complex dendritic arbors of the PVD nociceptive neurons (Inberg et al., 2019). The more penetrant dendritic phenotype of *grdn-1* mutants in AQR suggests that in the absence of interaction with glia or other surrounding cells, GRDN-1-dependent pathways may play a more critical role in regulating dendrite outgrowth in this neuron type.

Girdin domain organization is highly conserved, and this protein is considered to be a hub for multiple signal transduction pathways in different cell types (Aznar et al., 2016; Garcia-Marcos et al., 2015). We speculate that in *C. elegans*, GRDN-1 may similarly integrate multiple extrinsic and intrinsic signaling cues and couple them to diverse downstream effectors in order to regulate distinct phenotypes in a cell type-specific manner. Results from this work may guide future investigations into the underlying molecular mechanisms, and provide additional insights into the diverse roles of this multifunctional protein in different cellular contexts.

## Acknowledgements

We are grateful to Max Heiman, Guangshuo Ou, and Oliver Hobert for sharing reagents and advice, the *Caenorhabditis* Genetics Center for strains, and Max Heiman for discussion and sharing data prior to publication. We thank Anna Kazatskaya and Alison Philbrook for comments on the manuscript, and additional members of the Sengupta lab for advice. This work was funded in part by the National Institutes of Health (R35 GM122463 – P.S.) and startup funds from the Worcester Polytechnic Institute (I.V.N.).

## REFERENCES

Aznar, N., Kalogriopoulos, N., Midde, K.K., and Ghosh, P. (2016). Heterotrimeric G protein signaling via GIV/Girdin: Breaking the rules of engagement, space, and time. Bioessays 38, 379–393.

Bentley, M., and Banker, G. (2016). The cellular mechanisms that maintain neuronal polarity. Nat Rev Neurosci 17, 611–622.

Berger-Muller, S., and Suzuki, T. (2011). Seven-pass transmembrane cadherins: roles and emerging mechanisms in axonal and dendritic patterning. Mol Neurobiol 44, 313–320.

Byrd, D.T., Kawasaki, M., Walcoff, M., Hisamoto, N., Matsumoto, K., and Jin, Y. (2001). UNC-16, a JNK-signaling scaffold protein, regulates vesicle transport in *C. elegans*. Neuron 32, 787–800.

Cebul, E.R., McLachlan, I.G., and Heiman, M.G. (2020). Dendrites with specialized glial attachments develop by retrograde extension using SAX-7 and GRDN-1. Development 147, dev180448.

Chai, Y., Li, W., Feng, G., Yang, Y., Wang, X., and Ou, G. (2012). Live imaging of cellular dynamics during *Caenorhabditis elegans* postembryonic development. Nat Protoc 7, 2090–2102.

Costa, M., Raich, W., Agbunag, C., Leung, B., Hardin, J., and Priess, J.R. (1998). A putative catenin-cadherin system mediates morphogenesis of the *Caenorhabditis elegans* embryo. J Cell Biol 141, 297–308.

Dawe, H.R., Farr, H., and Gull, K. (2007). Centriole/basal body morphogenesis and migration during ciliogenesis in animal cells. J Cell Sci 120, 7–15.

Dokshin, G.A., Ghanta, K.S., Piscopo, K.M., and Mello, C.C. (2018). Robust genome editing with short single-stranded and long, partially single-stranded DNA donors in *Caenorhabditis elegans*. Genetics 210, 781–787.

Dong, X., Liu, O.W., Howell, A.S., and Shen, K. (2013). An extracellular adhesion molecule complex patterns dendritic branching and morphogenesis. Cell 155, 296–307.

Doroquez, D.B., Berciu, C., Anderson, J.R., Sengupta, P., and Nicastro, D. (2014). A high-resolution morphological and ultrastructural map of anterior sensory cilia and glia in *C. elegans*. eLife 3, e01948.

Elric, J., and Etienne-Manneville, S. (2014). Centrosome positioning in polarized cells: common themes and variations. Exp Cell Res 328, 240–248.

Enomoto, A., Ping, J., and Takahashi, M. (2006). Girdin, a novel actin-binding protein, and its family of proteins possess versatile functions in the Akt and Wnt signaling pathways. Ann NY Acad Sci 1086, 169–184.

Fan, L., Kovacevic, I., Heiman, M.G., and Bao, Z. (2019). A multicellular rosette-mediated collective dendrite extension. Elife 8, e38065.

Flavell, S.W., Pokala, N., Macosko, E.Z., Albrecht, D.R., Larsch, J., and Bargmann, C.I. (2013). Serotonin and the neuropeptide PDF initiate and extend opposing behavioral states in *C. elegans*. Cell 154, 1023–1035.

Garcia-Marcos, M., Ghosh, P., and Farquhar, M.G. (2015). GIV/Girdin transmits signals from multiple receptors by triggering trimeric G protein activation. J Biol Chem 290, 6697–6704.

Grana, T.M., Cox, E.A., Lynch, A.M., and Hardin, J. (2010). SAX-7/L1CAM and HMR-1/cadherin function redundantly in blastomere compaction and non-muscle myosin accumulation during *Caenorhabditis elegans* gastrulation. Dev Biol 344, 731–744.

Gray, J.M., Karow, D.S., Lu, H., Chang, A.J., Chang, J.S., Ellis, R.E., Marletta, M.A., and Bargmann, C.I. (2004). Oxygen sensation and social feeding mediated by a *C. elegans* guanylate cyclase homologue. Nature 430, 317–322.

Hall, D.H., and Russell, R.L. (1991). The posterior nervous system of the nematode *Caenorhabditis elegans:* serial reconstruction of identified neurons and complete pattern of synaptic interactions. J Neurosci 11, 1–22.

Harterink, M., Edwards, S.L., de Haan, B., Yau, K.W., van den Heuvel, S., Kapitein, L.C., Miller, K.G., and Hoogenraad, C.C. (2018). Local microtubule organization promotes cargo transport in *C. elegans* dendrites. J Cell Sci 131, jcs223107.

Heiman, M.G., and Shaham, S. (2009). DEX-1 and DYF-7 establish sensory dendrite length by anchoring dendritic tips during cell migration. Cell 137, 344–355.

Herman, M.A., and Horvitz, H.R. (1994). The *Caenorhabditis elegans* gene *lin-44* controls the polarity of asymmetric cell divisions. Development 120, 1035–1047.

Herman, M.A., Vassilieva, L.L., Horvitz, H.R., Shaw, J.E., and Herman, R.K. (1995). The *C. elegans* gene *lin-44*, which controls the polarity of certain asymmetric cell divisions, encodes a Wnt protein and acts cell nonautonomously. Cell 83, 101–110.

Holland, A.J., Fachinetti, D., Han, J.S., and Cleveland, D.W. (2012). Inducible, reversible system for the rapid and complete degradation of proteins in mammalian cells. Proc Natl Acad Sci USA 109, E3350–3357.

Houssin, E., Tepass, U., and Laprise, P. (2015). Girdin-mediated interactions between cadherin and the actin cytoskeleton are required for epithelial morphogenesis in *Drosophila*. Development 142, 1777–1784.

Ichimiya, H., Maeda, K., Enomoto, A., Weng, L., Takahashi, M., and Murohara, T. (2015). Girdin/GIV regulates transendothelial permeability by controlling VE-cadherin trafficking through the small GTPase, R-Ras. Biochem Biophys Res Commun 461, 260–267.

Inberg, S., Meledin, A., Kravtsov, V., Iosilevskii, Y., Oren-Suissa, M., and Podbilewicz, B. (2019). Lessons from worm dendritic patterning. Annu Rev Neurosci 42, 365–383.

Kazatskaya, A., Kuhns, S., Lambacher, N.J., Kennedy, J.E., Brear, A.G., McManus, G.J., Sengupta, P., and Blacque, O.E. (2017). Primary cilium formation and ciliary protein trafficking is regulated by the atypical MAP kinase MAPK15 in *Caenorhabditis elegans* and human cells. Genetics 207, 1423–1440.

Keil, T.A. (1997). Functional morphology of insect mechanoreceptors. Microsc Res Tech 39, 506–531.

Kirszenblat, L., Pattabiraman, D., and Hilliard, M.A. (2011). LIN-44/Wnt directs dendrite outgrowth through LIN-17/Frizzled in *C. elegans* neurons. PLoS Biol 9, e1001157.

Lefebvre, J.L. (2017). Neuronal territory formation by the atypical cadherins and clustered protocadherins. Semin Cell Dev Biol 69, 111–121.

Li, W., Yi, P., Zhu, Z., Zhang, X., Li, W., and Ou, G. (2017). Centriole translocation and degeneration during ciliogenesis in *Caenorhabditis elegans* neurons. EMBO J 36, 2553–2566.

Low, I.I.C., Williams, C.R., Chong, M.K., McLachlan, I.G., Wierbowski, B.M., Kolotuev, I., and Heiman, M.G. (2019). Morphogenesis of neurons and glia within an epithelium. Development 146.

Marston, D.J., Higgins, C.D., Peters, K.A., Cupp, T.D., Dickinson, D.J., Pani, A.M., Moore, R.P., Cox, A.H., Kiehart, D.P., and Goldstein, B. (2016). MRCK-1 drives apical constriction in *C. elegans* by linking developmental patterning to force generation. Curr Biol 26, 2079–2089.

McEwen, D.P., Jenkins, P.M., and Martens, J.R. (2008). Olfactory cilia: our direct neuronal connection to the external world. Curr Top Dev Biol 85, 333–370.

McLachlan, I.G., and Heiman, M.G. (2013). Shaping dendrites with machinery borrowed from epithelia. Curr Opin Neurobiol 23, 1005–1010.

Menco, B.P. (1997). Ultrastructural aspects of olfactory signaling. Chem Senses 22, 295–311.

Middelkoop, T.C., and Korswagen, H.C. (2014). Development and migration of the *C. elegans* Q neuroblasts and their descendants. WormBook, 1–23.

Muramatsu, A., Enomoto, A., Kato, T., Weng, L., Kuroda, K., Asai, N., Asai, M., Mii, S., and Takahashi, M. (2015). Potential involvement of kinesin-1 in the regulation of subcellular localization of Girdin. Biochem Biophys Res Commun 463, 999–1005.

Nechipurenko, I.V., Berciu, C., Sengupta, P., and Nicastro, D. (2017). Centriolar remodeling underlies basal body maturation during ciliogenesis in *Caenorhabditis elegans*. Elife 6, e25686.

Nechipurenko, I.V., Olivier-Mason, A., Kazatskaya, A., Kennedy, J., McLachlan, I.G., Heiman, M.G., Blacque, O.E., and Sengupta, P. (2016). A conserved role for Girdin in basal body positioning and ciliogenesis. Dev Cell 38, 493–506.

Nishimura, K., Fukagawa, T., Takisawa, H., Kakimoto, T., and Kanemaki, M. (2009). An auxin-based degron system for the rapid depletion of proteins in nonplant cells. Nat Methods 6, 917–922.

Oshita, A., Kishida, S., Kobayashi, H., Michiue, T., Asahara, T., Asashima, M., and Kikuchi, A. (2003). Identification and characterization of a novel Dvl-binding protein that suppresses Wnt signalling pathway. Genes Cells 8, 1005–1017.

Patel, N., Thierry-Mieg, D., and Mancillas, J.R. (1993). Cloning by insertional mutagenesis of a cDNA encoding *Caenorhabditis elegans* kinesin heavy chain. Proc Natl Acad Sci USA 90, 9181–9185.

Perkins, L.A., Hedgecock, E.M., Thomson, J.N., and Culotti, J.G. (1986). Mutant sensory cilia in the nematode *Caenorhabditis elegans*. Dev Biol 117, 456–487.

Rohlich, P. (1975). The sensory cilium of retinal rods is analogous to the transitional zone of motile cilia. Cell Tissue Res 161, 421–430.

Salzberg, Y., Diaz-Balzac, C.A., Ramirez-Suarez, N.J., Attreed, M., Tecle, E., Desbois, M., Kaprielian, Z., and Bulow, H.E. (2013). Skin-derived cues control arborization of sensory dendrites in *Caenorhabditis elegans*. Cell 155, 308–320.

Schafer, J.C., Haycraft, C.J., Thomas, J.H., Yoder, B.K., and Swoboda, P. (2003). XBX-1 encodes a dynein light intermediate chain required for retrograde intraflagellar transport and cilia assembly in *Caenorhabditis elegans*. Mol Biol Cell 14, 2057–2070.

Seong, E., Yuan, L., and Arikkath, J. (2015). Cadherins and catenins in dendrite and synapse morphogenesis. Cell Adh Migr 9, 202–213.

Serwas, D., Su, T.Y., Roessler, M., Wang, S., and Dammermann, A. (2017). Centrioles initiate cilia assembly but are dispensable for maturation and maintenance in *C. elegans*. J Cell Biol 216, 1659–1671.

Steimel, A., Wong, L., Najarro, E.H., Ackley, B.D., Garriga, G., and Hutter, H. (2010). The Flamingo ortholog FMI-1 controls pioneer-dependent navigation of follower axons in *C. elegans*. Development 137, 3663–3673.

Sulston, J.E., and Horvitz, H.R. (1977). Post-embryonic cell lineages of the nematode, *Caenorhabditis elegans*. Dev Biol 56, 110–156.

Sulston, J.E., Schierenberg, E., White, J.G., and Thomson, J.N. (1983). The embryonic cell lineage of the nematode *Caenorhabditis elegans*. Dev Biol 100, 64–119.

Takano, T., Xu, C., Funahashi, Y., Namba, T., and Kaibuchi, K. (2015). Neuronal polarization. Development 142, 2088–2093.

Tang, N., and Marshall, W.F. (2012). Centrosome positioning in vertebrate development. J Cell Sci 125, 4951–4961.

Wang, X., Enomoto, A., Weng, L., Mizutani, Y., Abudureyimu, S., Esaki, N., Tsuyuki, Y., Chen, C., Mii, S., Asai, N., et al. (2018). Girdin/GIV regulates collective cancer cell migration by controlling cell adhesion and cytoskeletal organization. Cancer Sci 109, 3643–3656.

Wang, X., Kweon, J., Larson, S., and Chen, L. (2005). A role for the *C. elegans* L1CAM homologue *lad-1/sax-7* in maintaining tissue attachment. Dev Biol 284, 273–291.

Ward, S., Thomson, N., White, J.G., and Brenner, S. (1975). Electron microscopical reconstruction of the anterior sensory anatomy of the nematode *Caenorhabditis elegans*. J Comp Neurol 160, 313–337.

Weng, L., Enomoto, A., Miyoshi, H., Takahashi, K., Asai, N., Morone, N., Jiang, P., An, J., Kato, T., Kuroda, K., et al. (2014). Regulation of cargo-selective endocytosis by dynamin 2 GTPase-activating protein girdin. EMBO J 33, 2098–2112.

White, J.G., Southgate, E., Thomson, J.N., and Brenner, S. (1986). The structure of the nervous system of the nematode *Caenorhabditis elegans*. Phil Transact R Soc Lond B 314, 1–340.

Yan, J., Chao, D.L., Toba, S., Koyasako, K., Yasunaga, T., Hirotsune, S., and Shen, K. (2013). Kinesin-1 regulates dendrite microtubule polarity in *Caenorhabditis elegans*. Elife 2, e00133.

Yogev, S., and Shen, K. (2017). Establishing neuronal polarity with environmental and intrinsic mechanisms. Neuron 96, 638–650.

Yu, S., Avery, L., Baude, E., and Garbers, D.A. (1997). Guanylyl cyclase expression in specific sensory neurons: A new family of chemosensory receptors. Proc Natl Acad Sci USA 94, 3384–3387.

Zhang, L., Ward, J.D., Cheng, Z., and Dernburg, A.F. (2015). The auxin-inducible degradation (AID) system enables versatile conditional protein depletion in *C. elegans*. Development 142, 4374–4384.

