## Supplementary data for "Girdin regulates dendrite morphogenesis and cilium position in two specialized sensory neuron types in *C. elegans*"

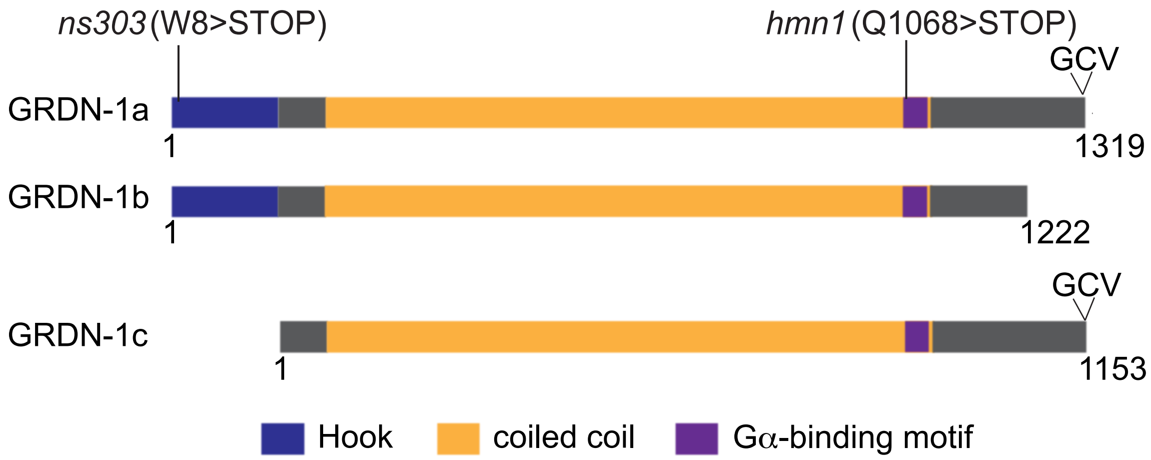


**Figure S1**

**Nechipurenko *et al.***

**Fig. S1**. Predicted isoforms of GRDN-1.

Conserved protein domains and premature termination codons in *grdn-1* alleles are shown. GCV indicates the predicted PDZ-binding motif.

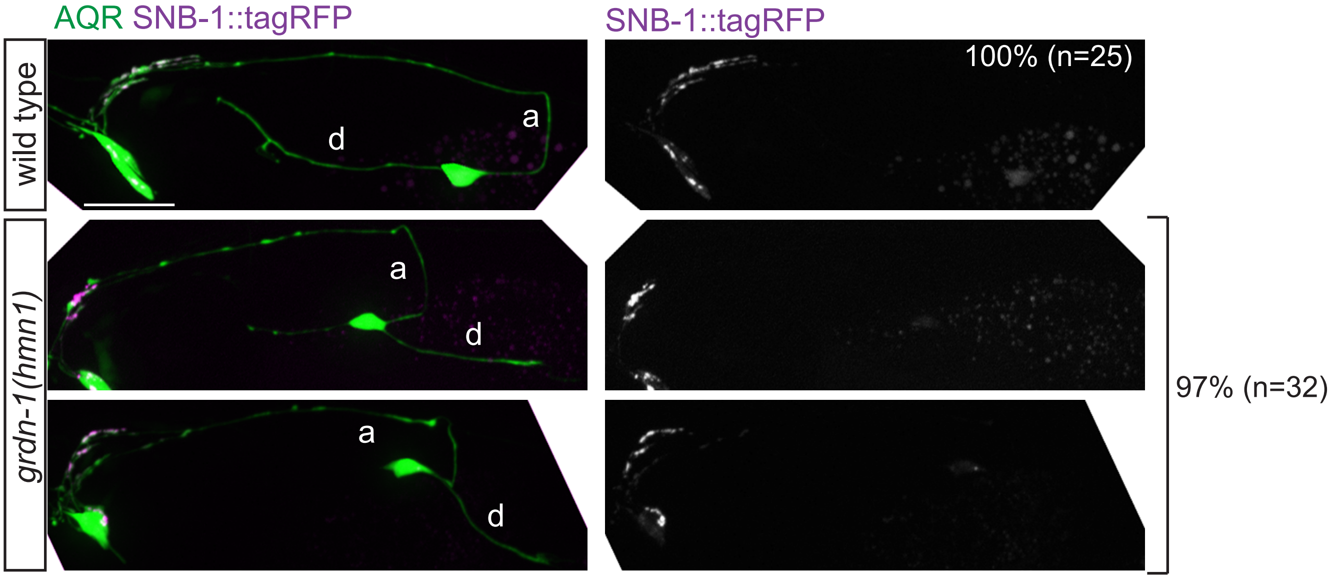


**Figure S2** **Nechipurenko *et al.***

**Fig. S2**. Localization of the synaptic protein SNB-1 is unaffected in AQR in *grdn-1* mutants. Images showing localization of *gcy-32*p::SNB-1::tagRFP in adult AQR neurons of wild-type and *grdn-1* mutant animals. AQR was visualized with *gcy-37*p::GFP. Percent of animals showing the depicted SNB-1::tagRFP localization pattern in AQR is indicated. a – axon; d – dendrite; anterior is at left. Scale bar: 10 μm.

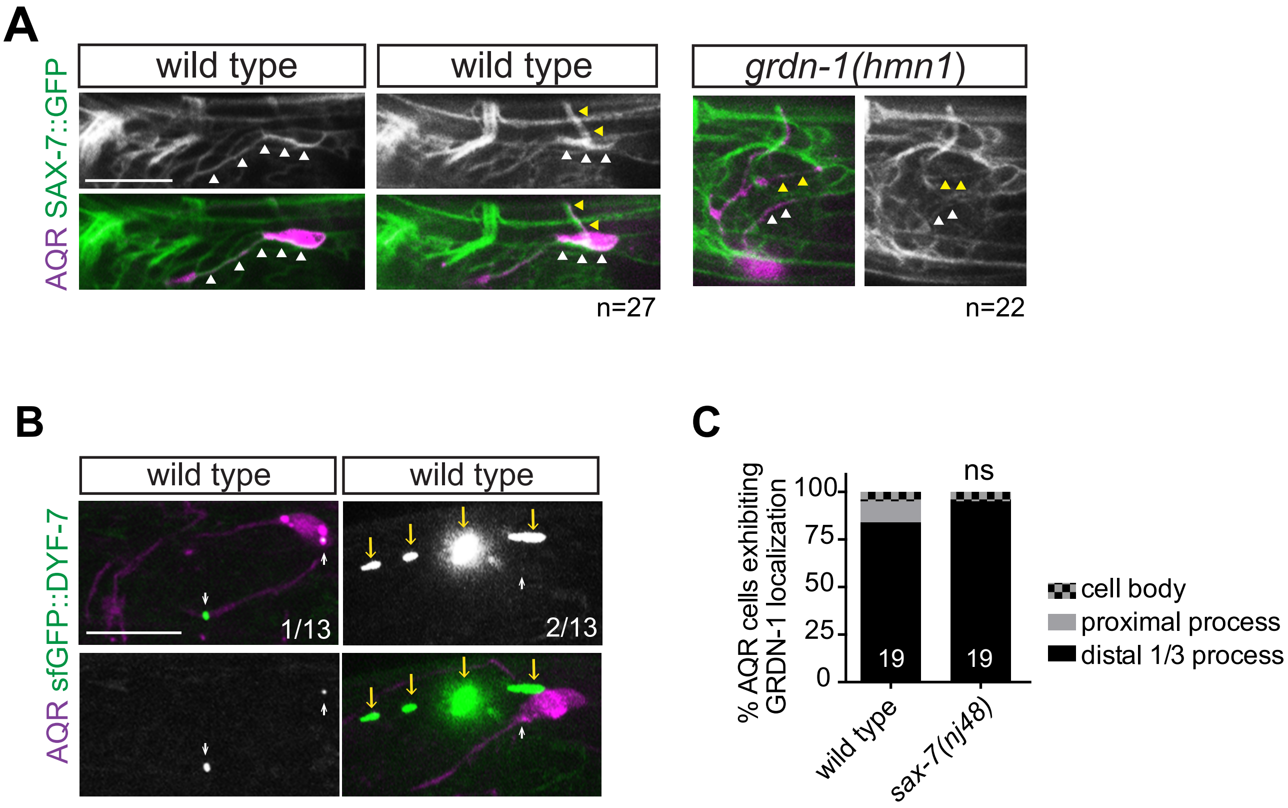


**Figure S3** Nechipurenko *et al.*

**Fig. S3**. Localization and expression of SAX-7 and DYF-7 in AQR neurons.

**(A)** A subset of optical sections from a confocal *z*-stack showing localization of the endogenously tagged SAX-7::GFP in AQR of wild-type and *grdn-1* mutant larvae. AQR was labeled with *egl-17*p::myr-mCherry. White arrowheads mark dendrites and cell bodies; yellow arrowheads mark axons. In all image panels, anterior is at left; scale bars: 5 μm.

**(B)** Images of *dyf-7*p::DYF-7::sfGFP expression in AQR in wild-type larvae. White arrows point to DYF-7::sfGFP signal in AQR. AQR was labeled with *egl-17*p::myr-mCherry. Proportion of animals with the depicted DYF-7::sfGFP localization pattern is indicated. sfGFP – superfolder GFP.

**(C)** Quantification of *gcy-36*p::GRDN-1::GFP localization in AQR in wild-type and *sax-7* animals. Numbers indicate number of AQR neurons examined per genotype. ns – not significantly different from wild-type (Fisher's exact test).

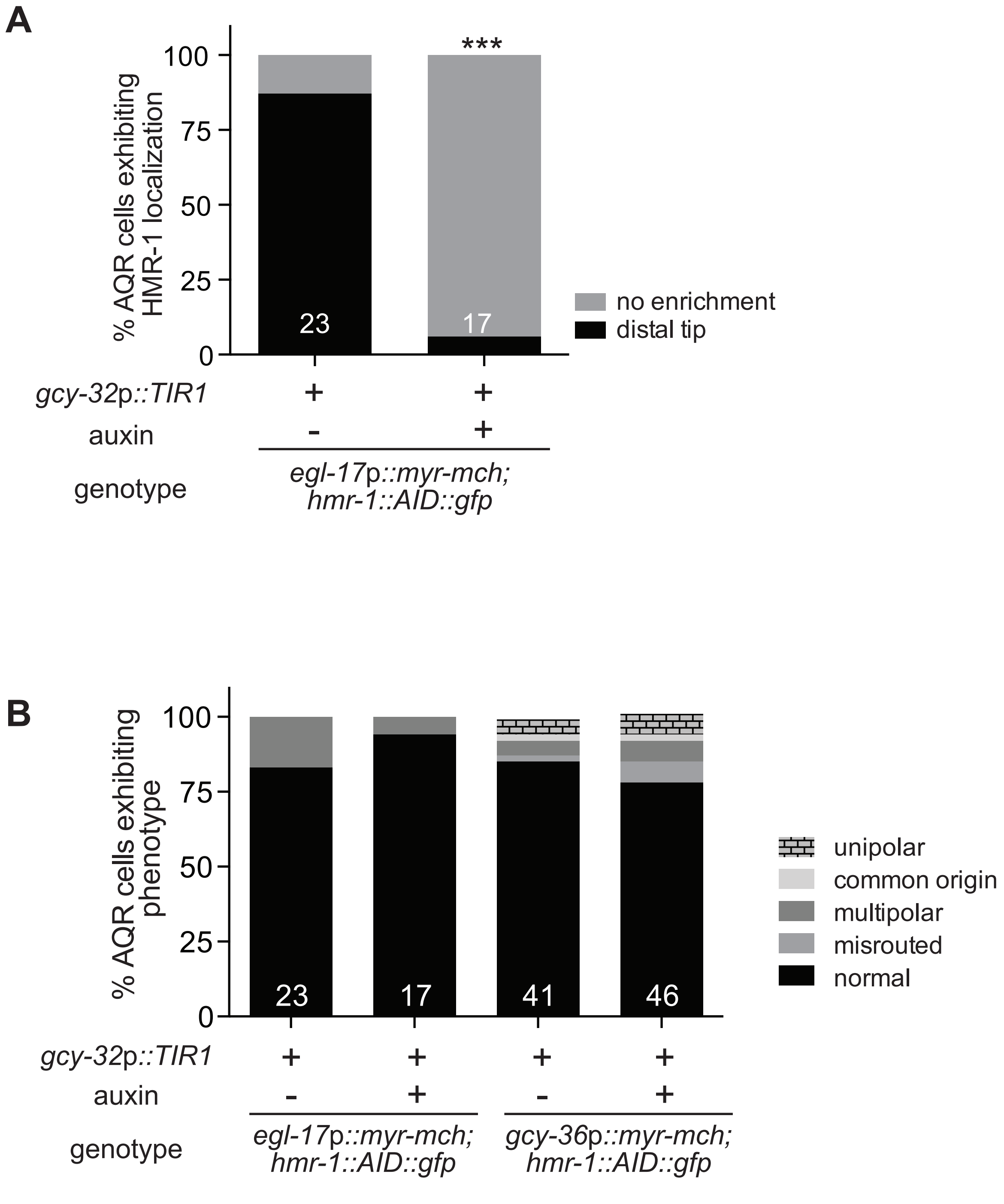
A
% AQR cells exhibiting HMR-1 localization
0
25
50
75
100
***
23
17
no enrichment
distal tip
gcy-32p::TIR1
+
+
auxin
-
+
genotype
egl-17p::myr-mch; hmr-1::AID::gfp
B
% AQR cells exhibiting phenotype
0
25
50
75
100
23
17
41
46
unipolar
common origin
multipolar
misrouted
normal
gcy-32p::TIR1
+
+
+
+
auxin
-
+
-
+
genotype
egl-17p::myr-mch; hmr-1::AID::gfp
gcy-36p::myr-mch; hmr-1::AID::gfp

**Figure S4**

**Fig. S4**. HMR-1 knockdown in AQR does not disrupt neuronal morphology.

Quantification of HMR-1::GFP signal **(A)** and AQR morphology **(B)** in the indicated experimental conditions. Numbers indicate number of AQR neurons examined per genotype. *** indicates different from wild-type at $p<0.001$ (Fisher's exact test).

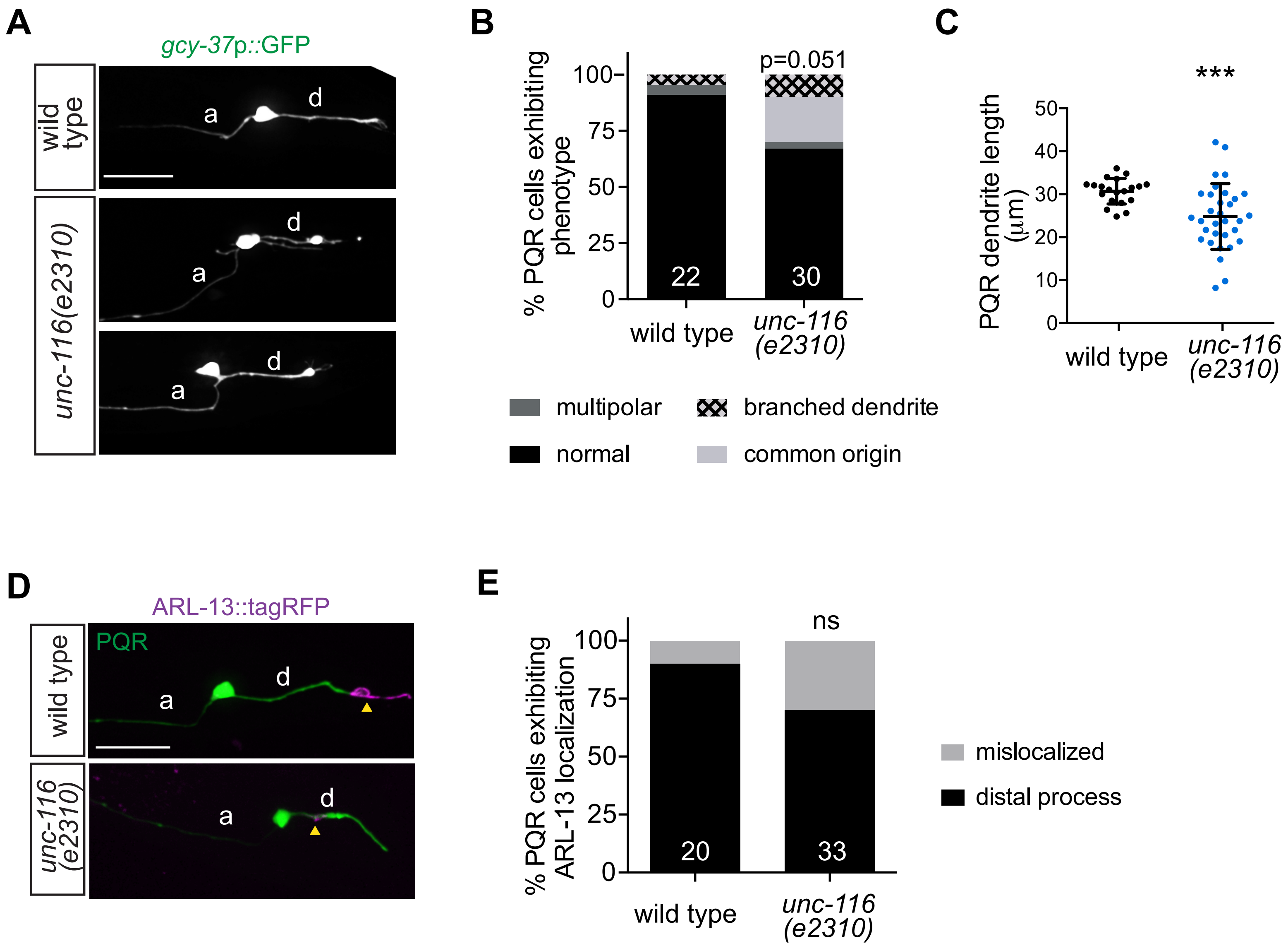


**Figure S5** **Nechipurenko *et al.***

**Fig. S5**. PQR morphology and cilium position are largely unaffected in *unc-116* animals.

**(A-B)** Images **(A)** and quantification **(B)** of PQR morphologies in adult wild-type and *unc-116* mutant animals. p-value in (**B**) derived from Fisher's exact test.

**(C)** Quantification of dendrite length in adult wild-type and *unc-116* mutant animals. *** indicates different from wild-type at $p<0.001$ (Student t test with Welch's correction).

**(D–E)** Images **(D)** and quantification **(E)** of *gcy-32*p::ARL-13::tagRFP localization in PQR of wild-type and *unc-116* mutant animals. PQR was visualized with *gcy-37*p::GFP. ns – not significantly different from wild type (Fisher's exact test).

In all image panels, a – axon; d – dendrite; anterior is at left; scale bars: 10 μm. In all bar graphs, numbers indicate number of PQR neurons examined per genotype.

**Table S1.** Strains used in this work

| Strain | Genotype | Source |
|---|---|---|
| PY11316 | *ials25[gcy-37*p*::gfp, unc-119(+)]; oyEx645[gcy-32*p*::arl-13::tagRfp, unc-122*p*::mCherry]* | This work |
| PY11318 | *oyEx640[grdn-1*p*::grdn-1a::gfp, gcy-32*p*::mCherry]* | This work |
| PY11361 | *oyEx641[gcy-32*p*::arl-13::tagRfp, gcy-36*p*::grdn-1a::gfp, unc-122*p*::dsRed]* | This work |
| PY11379 | *oyEx642[gcy-32*p*::xbx-1::tagRfp, gcy-36*p*::grdn-1a::gfp, unc-122*p*::dsRed]* | This work |
| PY11327 | *ials25[gcy-37*p*::gfp, unc-119(+)]; oyEx643[gcy-36*p*::dyf-19::tagRfp, unc-122*p*::mCherry]* | This work |
| PY11340 | *casIs165[egl-17*p*::myr-mCherry, pRF4(+)]; oyEx644[gcy-36*p*::grdn-1a::gfp, srg-47*p*::gfp]* | This work |
| PY11326 | *ials25[gcy-37*p*::gfp, unc-119(+)]; grdn-1(ns303); oyEx643[gcy-36*p*::dyf-19::tagRfp, unc-122*p*::mCherry]* | This work |
| PY11350 | *ials25[gcy-37*p*::gfp, unc-119(+)]; grdn-1(hmn1); oyEx643[gcy-36*p*::dyf-19::tagRfp, unc-122*p*::mCherry]* | This work |
| PY11342 | *ials25[gcy-37*p*::gfp, unc-119(+)]; grdn-1(ns303); oyEx645[gcy-32*p*::arl-13::tagRfp, unc-122*p*::mCherry]* | This work |
| PY11317 | *ials25[gcy-37*p*::gfp, unc-119(+)]; grdn-1(hmn1); oyEx645[gcy-32*p*::arl-13::tagRfp, unc-122*p*::mCherry]* | This work |
| PY11330 | *ials25[gcy-37*p*::gfp, unc-119(+)]; grdn-1(hmn1); oyEx646[grdn-1*p*::grdn-1a, unc-122*p*::dsRed]; oyEx647[gcy-32*p*::arl-13::tagRfp, unc-122*p*::gfp]* | This work |
| PY11359 | *ials25[gcy-37*p*::gfp, unc-119(+)]; grdn-1(hmn1); oyEx645[gcy-32*p*::arl-13::tagRfp, unc-122*p*::mCherry]; oyEx669[rab-3*p*::grdn-1DGCV, unc-122*p*::gfp]* | This work |
| ZG610 | *iaIs25[gcy-37*p*::gfp, unc-119(+)]* | CGC |
| PY9445 | *iaIs25[gcy-37*p*::gfp, unc-119(+)]; grdn-1(ns303)* | This work |
| PY11341 | *[gcy-37*p*::gfp, unc-119(+)]; grdn-1(hmn1)* | This work |
| PY11376 | *ials25[gcy-37*p*::gfp, unc-119(+)]; grdn-1(hmn1); oyEx653[grdn-1*p*::grdn-1a, unc-122*p*::dsRed]* | This work |
| PY11378 | *ials25[gcy-37*p*::gfp, unc-119(+)]; grdn-1(hmn1); oyEx646[grdn-1*p*::grdn-1a, unc-122*p*::dsRed]* | This work |
| PY11344 | *iaIs25[gcy-37*p*::gfp, unc-119(+)]; grdn-1(hmn1); oyEx670[grdn-1*p*::grdn-1DGCV, unc-122*p*::mCherry]* | This work |
| PY11321 (derived from GOU812) | *casIs165[egl-17*p*::myr-mCherry,* pRF4(+)*]* | A gift of G. Ou (Tsinghua University, PRC) |
| PY11322 | *casIs165[egl-17*p*::myr-mCherry*, pRF4(+)*]; grdn-1(hmn1)* | This work |
| PY11352 | *casIs165[egl-17*p*::myr-mCherry*, pRF4(+)*]; grdn-1(hmn1); oyEx650[rab-3*p*::grdn-1::gfp, srg-47*p*::gfp]* | This work |
| PY11364 | *casIs165[egl-17*p*::myr-mCherry,* pRF4(+)*]; grdn-1(hmn1); oyEx652[rab-3*p*::grdn-1DGCV, unc-122*p*::gfp]* | This work |

| | | |
|---|---|---|
| PY11377 | *casIs165[egl-17*p*::myr-mCherry,* pRF4(+)*]; grdn-1(hmn1); oyEx651[rab-3*p*::grdn-1DGCV, unc-122*p*::gfp]* | This work |
| PY11337 | *ials25[gcy-37*p*::gfp, unc-119(+)]; oyEx648[gcy-32*p*::snb-1::tagRfp, unc-122*p*::gfp]* | This work |
| PY11338 | *ials25[gcy-37*p*::gfp, unc-119(+)]; grdn-1(hmn1); oyEx648[gcy-32*p*::snb-1::tagRfp, unc-122*p*::gfp]* | This work |
| PY11324 | *cp21[hmr-1::gfp + LoxP]; casIs165[egl-17*p*::myr-mCherry,* pRF4(+)*]* | This work |
| PY11355 | *casIs165[egl-17*p*::myr-mCherry,* pRF4(+)*]; ddIs290[sax-7::TY1::egfp::3xflag(92C12) + unc-119(+)]* | This work |
| PY11380 | *iaIs25[egl-17*p*::myr-mCherry,* pRF4(+)*]; hmnEx146[dyf-7*p*::sfgfp::dyf-7,* pRF4(+)] | This work |
| PY11349 | *casIs165[egl-17*p*::myr-mCherry,* pRF4(+)*]; dlg-1(cp301[dlg-1::mNG-C1^3xflag])* | This work |
| PY11346 | *cp21[hmr-1::gfp + LoxP]; casIs165[egl-17*p*::myr-mCherry,* pRF4(+)]; *grdn-1(ns303)* | This work |
| PY11325 | *cp21[hmr-1::gfp + LoxP]; casIs165[egl-17*p*::myr-mCherry,* pRF4(+)*]; grdn-1(hmn1)* | This work |
| PY11371 | *casIs165[egl-17*p*::myr-mCherry*, pRF4(+)*]; ddIs290 [sax-7::TY1::egfp::3xflag(92C12) + unc-119(+)]; grdn-1(hmn1)* | This work |
| PY11370 | *casIs165[egl-17*p*::myr-mCherry*, pRF4(+)*]; sax-7(nj48); oyEx644[gcy-36*p*::grdn-1a::gfp, srg-47*p*::gfp]* | This work |
| PY11347 | *hmr-1(hd37); ials25[gcy-37*p*::gfp, unc-119(+)]; oyEx645[gcy-32*p*::arl-13::tagRfp, unc-122*p*::mCherry]* | This work |
| PY11372 | *hmr-1(oy157[hmr-1::AID+gfp]); oyIs89[gcy-36*p*::myr-tagRfp]; oyEx656[gcy-32*p*::TIR1, unc-122*p*::dsRed]* | This work |
| PY11368 | *hmr-1(oy157[hmr-1::AID+gfp]); casIs165[egl-17*p*::myr-mCherry,* pRF4(+)*]; oyEx655[gcy-32*p*::TIR1, unc-122*p*::dsRed]* | This work |
| PY11348 | *ials25[gcy-37*p*::gfp, unc-119(+)]; sax-7(ky146); oyEx645[gcy-32*p*::arl-13::tagRfp, unc-122*p*::mCherry]* | This work |
| PY11360 | *ials25[gcy-37*p*::gfp, unc-119(+)]; sax-7(eq1); oyEx645[gcy-32*p*::arl-13::tagRfp, unc-122*p*::mCherry]* | This work |
| PY11359 | *ials25[gcy-37*p*::gfp, unc-119(+)]; sax-7(nj48); oyEx645[gcy-32*p*::arl-13::tagRfp, unc-122*p*::mCherry]* | This work |
| PY11374 | *hmr-1(hd37); ials25[gcy-37*p*::gfp, unc-119(+)]; sax-7(nj48); oyEx645[gcy-32*p*::arl-13::tagrfp, unc-122*p*::mCherry]* | This work |
| PY11335 | *unc-116(e2310); casIs165[egl-17*p*::myr-mCherry,* pRF4(+)*]; oyEx644[gcy-36*p*::grdn-1a::gfp, srg-47*p*::gfp]* | This work |
| PY11334 | *cp21[hmr-1::gfp + LoxP]; unc-116(e2310); casIs165[egl-17*p*::myr-mCherry,* pRF4(+)*]* | This work |
| PY11345 | *unc-116(e2310); ials25[gcy-37*p*::gfp, unc-119(+)]; oyEx645[gcy-32*p*::arl-13::tagRfp, unc-122*p*::mCherry]* | This work |
| PY11351 | *unc-116(ce815); casIs165[egl-17*p*::myr-mCherry;*pRF4(+)] | This work |
| PY11375 | *unc-116(ce815); casIs165[egl-17*p*::myr-mCherry,* pRF4(+)*]; oyEx654[tag-168*p*::Cre, unc-122*p*::gfp]* | This work |
| NWM001 | *lin-44(n1792); casIs165[egl-17*p*::myr-mCherry;*pRF4(+)*]; oyEx644[gcy-36*p*::grdn-1a::gfp, srg-47*p*::gfp]* | This work |
| NWM002 | *lin-44(n1792); ials25[gcy-37*p*::gfp, unc-119(+)]; oyEx645[gcy-32*p*::arl-13::tagRfp, unc-122*p*::mCherry]* | This work |

**Table S2.** Plasmids used in this work

| Plasmid | Description | Source |
|---|---|---|
| PSAB1002 | *grdn-1*p*::grdn-1a::gfp* | Nechipurenko *et al*., 2016 |
| PSAB1215 | *gcy-32*p*::arl-13::tagRfp* | This work |
| PSAB1217 | *gcy-36*p*::grdn-1a::gfp* | This work |
| PSAB1218 | *gcy-32*p*::xbx-1::tagRfp* | This work |
| PSAB1216 | *gcy-36*p*::dyf-19::tagRfp* | This work |
| PSAB988 | *grdn-1p::grdn-1a* | Nechipurenko *et al*., 2016 |
| PSAB1219 | *rab-3*p*::grdn-1*$^{\Delta GCV}$ | This work |
| PSAB1220 | *grdn-1*p*::grdn-1*$^{\Delta GCV}$ | This work |
| PSAB1221 | *rab-3*p*::grdn-1a::gfp* | This work |
| PSAB1013 | *srg-47*p*::gfp* | Nechipurenko *et al*., 2016 |
| PSAB1222 | *gcy-32*p*::snb-1::tagRfp* | This work |
| PSAB1223 | *gcy-32*p*::TIR1* | This work |
| pSF11 | *tag-168*p*::Cre (nCre)* | Flavell *et al*., 2013 |
